# Suppression of epithelial proliferation and tumourigenesis by IgA

**DOI:** 10.1101/2023.10.06.561290

**Authors:** Gregory P. Donaldson, Gabriella L. Reis, Marwa Saad, Christopher Wichmann, Izabel Mamede, Guo Chen, Caroline E. Harrer, Nicole L. DelGaudio, Dayu Zhang, Begüm Aydin, Tiago B.R. Castro, Sergei I. Grivennikov, Bernardo S. Reis, Elise G. Liu, Stephanie C. Eisenbarth, Kivanç Birsoy, Sohail F. Tavazoie, Beth M. Stadtmueller, Gabriel D. Victora, Daniel Mucida

**Author notes:** For materials and correspondence, contact: Gregory P. Donaldson or Daniel Mucida. Department of Microbiology, Immunology, & Molecular Genetics, University of California, Los Angeles, United States (Gregory P. Donaldson).

## Abstract

Immunoglobulin A (IgA) is the most abundant antibody isotype produced by mammals, playing a specialised role in mucosal homeostasis^1^. IgA is constantly secreted into the intestinal lumen and binds commensal microbiota to regulate their colonisation and function^2,3^, with unclear implications for health. IgA deficiency is common in humans but is difficult to study due to its complex aetiology and comorbidities^4–8^. Using genetically and environmentally controlled mice, here we show that multiple animal models of IgA deficiency result in increased susceptibility to multiple models of endogenous colorectal tumours. Cellular and molecular analyses revealed that, in the absence of IgA, colonic epithelial cells induce antibacterial factors and accelerate cell cycling in response to the microbiota. Oral treatment with IgA was sufficient to reduce steady-state proliferation and protect mice from tumours; however, this function was due to the antibody structure rather than binding specificity. In a reductionist culture system, IgA directly suppresses epithelial growth. Co-immunoprecipitation mass spectrometry and targeted CRISPR screening identified DMBT1 as an IgA-binding epithelial surface protein required for IgA-mediated suppression of proliferation. Together, IgA and DMBT1 tune the normal cycling of absorptive colonocyte progenitors. In mice, deletion of the transmembrane and cytoplasmic signalling portions of DMBT1 was sufficient to reverse both the increased proliferation and tumour susceptibility of IgA knockouts. These experiments establish a homeostatic function for IgA in tempering physiological epithelial responses to microbiota to maintain mucosal health.

---

In the absence of infection or disease, IgA is constantly produced in the intestine, where it is secreted into the lumen in the form of secretory IgA (SIgA) and binds commensal microbiota^9^. In humans, selective IgA deficiency is the most common primary immunodeficiency, clinically defined as low (<7 mg/dL) serum IgA levels with normal levels of IgM and IgG^10^. These patients have an increased risk of respiratory and gastrointestinal infections, inflammatory and autoimmune disorders^10^, and gastrointestinal cancers^11^. IgA is known to control mucosal pathogens^12–14^; however, its impact on inflammatory diseases and malignancy remains unclear. Recent studies have indicated that IgA deficiency is associated with changes in the microbiota and systemic inflammation^4,6,7,15^, which may be more pronounced in patients with undetectable IgA^8^. A clinical association with malignancy was observed decades ago^16,17^. The largest study on IgA deficiency (2,495 against 24,509 controls) showed double all-cause mortality and an increased risk of colon and prostate cancers^18^. Inverse analyses of patients with gastrointestinal cancers also found an increased frequency of IgA deficiency^19,20^. However, the polygenic nature of IgA deficiency, its arbitrary clinical definition based on serum concentration, and various comorbidities thwart the attribution of these health effects to isotype-specific functions.

Most known functions of intestinal IgA have been discovered using mouse models, showing that these antibodies prevent certain mucosal infections^12–14^ and affect the form and function of the gut microbiome^21–28^. Effects on bacteria vary; IgA controls *Bacteroides thetaiotaomicron* gene expression^29,30^, *Salmonella enterica* swimming motility^14^, *Bacteroides fragilis* mucosal anchoring^31^, and *Escherichia coli* surface receptor activities^32^. It remains unclear how these functions impact health. An isotype-specific knockout mouse model (*Igha*^−/−,^ hereafter referred to as IgA^−/−^) causing total IgA deficiency was established more than 20 years ago^33^. These animals have increased serum and mucosal IgM^33^, mirroring human IgA deficiency^4^. IgA^−/−^ mice have only a few described phenotypes^13^ and are largely indistinguishable from wild-type controls in a dextran sodium sulfate colitis model^34^, suggesting that IgA is dispensable for the normal intestinal barrier. In contrast, polymeric immunoglobulin receptor (pIgR) knockouts, which are unable to actively transport antibodies to the lumen of the intestine, are more susceptible to colitis^34^, highlighting the importance of compensatory IgM or other functions of pIgR in supporting the barrier. The antibody-binding ectodomain of pIgR, known as the secretory component, is secreted even in the absence of IgA and may have independent functions^35,36^. Despite evidence that it is dispensable in certain contexts, IgA is the most abundantly produced isotype in humans.

## IgA deficiency impacts cancer models

To study the direct physiological consequences of IgA deficiency, female IgA^+/−^ mice were bred with IgA^−/−^ males to control for effects of maternal milk IgA^37^. IgA^+/−^ mice had comparable levels of faecal IgA to IgA^+/+^ littermates (**Extended Data Fig. 1a-b**), confirming that heterozygotes can serve as normal controls. We observed compensatory induction of faecal IgG and IgM in IgA^−/−^ mice, but not to the normal level of faecal IgA (**Extended Data Fig. 1c-e**). Co-housed littermates were used in all experiments to minimise environmental variables and normalise the microbiome. For an unbiased analysis of possible IgA effects, mid-colon tissues were sampled at steady state for bulk RNA-seq (**Fig. 1a**). While the total number of differentially expressed genes was modest, IgA^−/−^ mice exhibited epithelial-specific changes, including a decrease in goblet cell markers (*Spink4*, *Fcgbp*, *Capn9,* and *Tff3*) and other epithelial cell subtype-specific markers, such as *SpiB* (M cell transcription factor) and *Rgs4* (enteroendocrine cell marker) (**Fig. 1a, Supplementary Data 1**). This coincided with an increase in *Nox1*, marking highly proliferative transit-amplifying cells^38^, with concordant increases in cell cycling-related pathways and genes (*Mcc* and *Sass6*) (**Fig. 1a**). These results indicate a physiological role for IgA in balancing proliferation and differentiation in the intestinal epithelium under steady-state conditions.

**Fig. 1.**
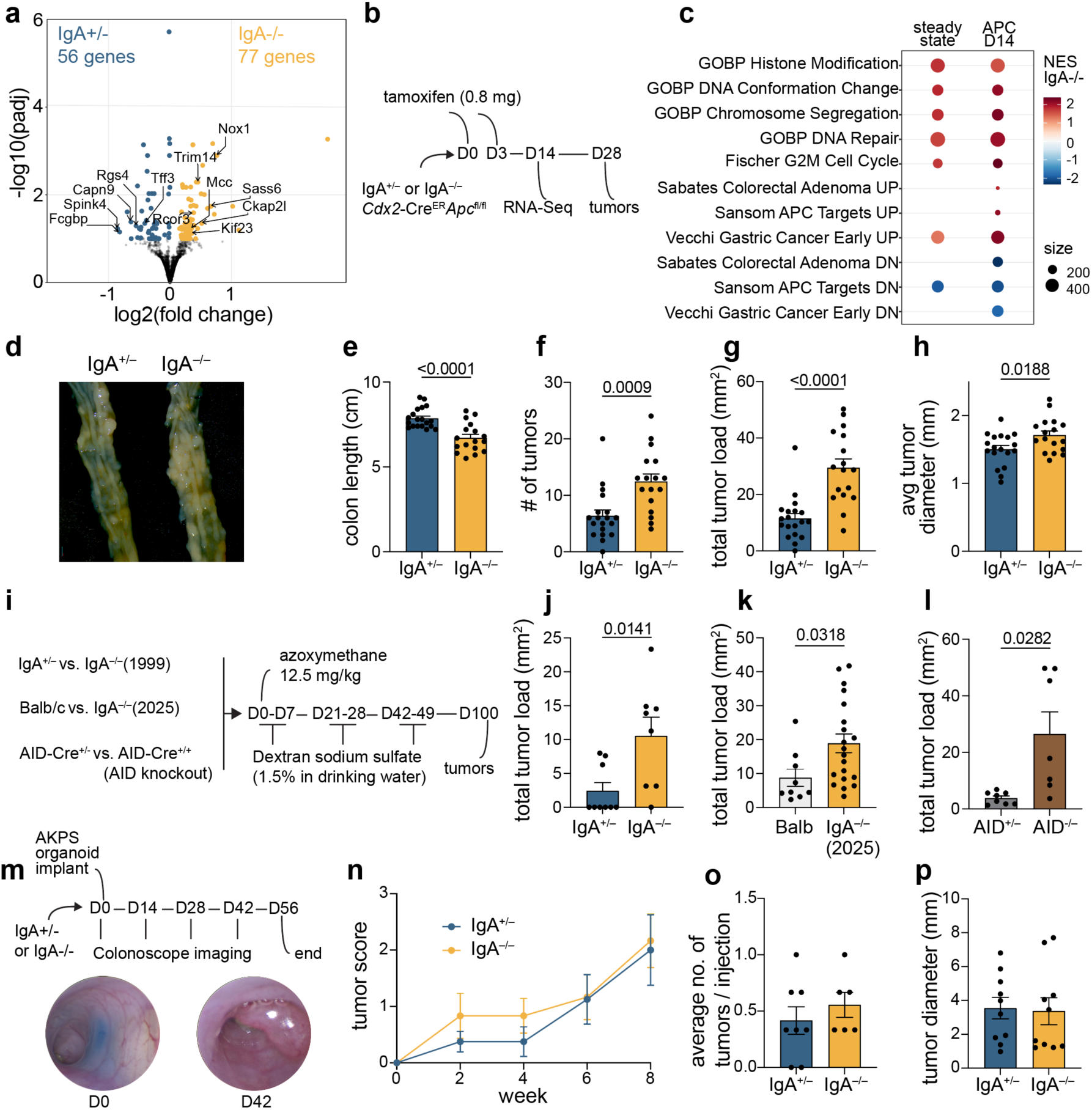
IgA^−/−^ mice have pro-tumour alterations to colon tissue and increased susceptibility to endogenous colorectal cancer models. **a,** Bulk RNA-Seq of colon tissue from IgA^+/−^ and IgA^−/−^ mice during steady state revealed upregulation of transit-amplifying cell markers (DESeq2, n = 5). b, Experimental design for studies of IgA in the context of epithelial-specific loss of *APC*. c, Gene set enrichment analyses indicated a pro-cancerous phenotype of IgA^−/−^ mice that was exacerbated after APC loss (n = 5 steady, 4 APC, visible dots only for gene sets where adjusted p < 0.05). d-h, On day 28, mice were sacrificed and assessed for colon length, number of colon tumours, total tumour load, and average tumour size (pooled, 4 cohorts, n = 19, 17). i, Experimental design for azoxymethane (AOM) and dextran sodium sulfate (DSS) model of colorectal cancer in 3 models of IgA deficiency. Mice were sacrificed at day 100 to assess tumour outcomes in j, 1999 IgA^−/−^ mice (pooled 2 cohorts, n = 9, 8), k, 2025 IgA^−/−^ mice (pooled 2 cohorts, n = 9, 20, mice aged 3-8 months, non-littermate controls from same room, due to limited availability), and l, AID^−/−^ mice (representative of 2 cohorts, n = 8, 7). m, Experimental design and example images for colonoscope-guided implant of quadruple-mutant (*APC*, *Kras*, *P53*, *Smad4*) organoid cells. n, Tumour growth tracked longitudinally by colonoscope and scored based on the size of the tumour relative to the colon circumference (pooled 2 cohorts, n = 8, 6, repeated- measure 2-way ANOVA not significant). o, After 8 weeks, mice were sacrificed to assess implantation rate per injection and p, tumour size (n = 10). (mean and SEM, two-tailed t-tests unless stated otherwise, p values hidden when p > 0.1)

Next, we used a murine model of endogenous colon cancer as a perturbation to test the effects of IgA in a relevant disease state. IgA^−/−^ mice were crossed with a strain harbouring a *Cdx2* promoter-driven, tamoxifen-inducible Cre recombinase and loxP-flanked alleles of the tumour suppressor *APC*, which is mutated in more than 90% of human colorectal tumours^39^. This allowed for the temporally controlled homozygous knockout of *APC* in a fraction of colonic epithelial cells (**Fig. 1b**). We harvested tissue for a parallel RNA-Seq at an early, pre-tumour time point (**Fig. 1b-c**). Despite synchronised loss of APC, there were substantial (hundreds of genes) differences in gene expression patterns between IgA^+/−^ and IgA^−/−^ animals (**Extended Data Fig. 1f, Supplementary Data 2**). Specifically, pathways related to epithelial cell proliferation, APC mutation, and human colorectal cancer were enhanced in IgA^−/−^ mice (**Fig. 1c**). At the pathway level, modest alterations at steady state in IgA^−/−^ mice mirrored the more dramatic phenotype induced by the APC mutation (**Fig. 1c**). Most genes altered during steady state by IgA genotype were also modulated by *APC* loss, with nearly perfect directional concordance between IgA deficiency and *APC* loss (**Extended Data Fig. 1g**). These molecular results were also predictive of tumour outcomes (**Fig. 1d**). While IgA^+/+^ and IgA^+/−^ mice were indistinguishable (**Extended Data Fig. 1h**), IgA^−/−^ mice had shortened colons (**Fig. 1e**), increased number of tumours (**Fig. 1f**), increased total tumour load (**Fig. 1g**), and slightly increased tumour size (**Fig. 1h**) compared to cohoused IgA^+/−^ littermates. Although this cancer model is dependent on inflammation^39^, faecal lipocalin as a measure of inflammation (**Extended Data Fig. 1i**), and expression levels of cytokines in bulk RNA-seq (**Extended Data Fig. 1j**) showed no differences in IgA^−/−^ mice. Flow cytometry analysis of adaptive and innate immune cells in the colon lamina propria and tumours did not reveal clear differences in the immune response to tumours in IgA^+/−^ vs. IgA^−/−^ littermates (**Extended Data Fig. 2**). These molecular and tumour analyses indicated that IgA^−/−^ mice are more susceptible to an *APC*-driven model of colorectal cancer, but this did not correlate with inflammation.

To expand to other models of colorectal cancer, IgA^+/−^ and IgA^−/−^ mice were subjected to injection of the mutagen azoxymethane, followed by three rounds of exposure to dextran sodium sulfate (AOM-DSS) in their drinking water (**Fig. 1i**), mimicking colitis-associated colorectal cancer. As previously reported^34^, IgA^−/−^ mice were slightly more susceptible to DSS-induced weight loss (**Extended Data Fig. 3a**). Using a relatively low dose of DSS (1.5%), most IgA^−/−^ mice developed tumours while most IgA^+/−^ mice developed none (**Extended Data Fig. 3b**). IgA^−/−^ mice showed higher overall tumour load and number of tumours compared to IgA^+/−^ controls (**Fig. 1j, Extended Data Fig. 3c**). We further tested other models of IgA deficiency, including a recently made IgA knockout that targeted the first exon of the alpha heavy chain^40^. Although in a different location (Yale) and different mouse genetic background (BALB/c), these mice also exhibited greater overall tumour load compared to wildtype control mice when subjected to the same AOM-DSS protocol (**Fig. 1k, Extended Data Fig. 3d**). Mice homozygous for a knock-in *Aid*-*Cre* allele are effectively knockouts of activation-induced cytidine deaminase, which is required for class switch recombination, resulting in B cell restriction to isotypes IgM or IgD. With the caveat that this also results in IgG deficiency, in the AOM-DSS model, AID knockout animals also had greater tumour load compared to heterozygote controls (**Fig. 1l**, **Extended Data Fig. 3e-f**). These results with two colorectal cancer models (that result in tumours in different colon locations which have physiological differences, **Extended Data Fig. 3g**) and three IgA deficiency models were generally consistent in revealing a protective role for IgA in the context of colorectal tumour formation.

Next, we tested the impact of IgA on an implant model, where orthotopic injection of cells results in tumour growth within the subepithelial dome, as opposed to the epithelial surface, as in endogenous models. Mouse colon organoids with four cancer-related gene alterations (knockout of *APC*, *Smad4*, and *P53,* plus activated *Kras* G12D) were dissociated into single cells, injected directly into the colon wall using a colonoscope, and monitored by colonoscopy (**Fig. 1m**). Regardless of the host IgA genotype, these implanted tumours grew at a similar rate (**Fig. 1n**), with a similar implantation success frequency (**Fig. 1o**), and were similar in size at the endpoint (**Fig. 1p**). Overall, our data indicate that IgA has a reproducible protective function in endogenous colorectal tumour models but not implant models with exogenous cells.

## IgA-microbiome interactions

As the primary known function of IgA at steady state is to regulate the composition and function of the microbiota, we investigated whether IgA^−/−^ mice harboured a pro-tumour microbiome despite being cohoused with littermates. The role of bacteria was first assessed by treating mice with a cocktail of broad-spectrum antibiotics (ampicillin, vancomycin, metronidazole, and neomycin) throughout the experiment (**Fig. 2a**), which did not affect faecal IgA levels (**Extended Data Fig. 4a**). This treatment was previously reported to reduce tumour numbers in this model^39^, and in the context of antibiotic depletion we found that IgA^−/−^ mice did not exhibit increased tumourigenesis (**Fig. 2b**), linking this phenomenon to the microbiota. Next, we profiled the microbiome using sequencing to determine whether alterations in the microbial composition might explain the observations in Fig. 1. 16S rRNA amplicon sequencing was performed in healthy mice at steady state and two weeks after *APC* deletion. Microbiomes from IgA^+/−^ and IgA^−/−^ mice were indistinguishable from those of littermate controls when co-housed throughout the experiment (**Fig. 2c, Extended Data Fig. 4b**). *APC* loss caused a shift in the microbiome, with a loss of diversity, including the depletion of species within the common mouse commensal family, *Muribaculaceae* (**Fig. 2c**). This coincided with the outgrowth of *Shigella* and *Helicobacter* species (**Fig. 2c**), both of which have been implicated in colorectal cancer^41^. During this dramatic change, IgA still had no apparent influence on microbiome composition during these experiments in which littermates were co-housed from birth (**Fig. 2c, Extended Data Fig. 4b**). As some IgA-mediated alterations have previously been shown to be mucus-specific^23,31^, colon mucus was aseptically sampled for microbiome profiling. This analysis successfully resolved the spatial differences in lumen and mucosal community compositions (**Extended Data Fig. 4=b-c**)^42^ and revealed that some commensal mucosal species of *Akkermansia* and *Bacteroides* were resilient to *APC* loss (**Extended Data Fig. 4c**). However, even in mucus, no differences were apparent between IgA^+/−^ and IgA^−/−^ samples (**Supplementary Fig. 1a**). The microbiome shift observed during *APC* loss was not observed in the AOM-DSS model (**Extended Data Fig. 4d, Supplementary Fig. 1b**). This difference was not due to the impact of tamoxifen (**Extended Data Fig. 4e, Supplementary Fig. 1c**), but perhaps due to tumours being restricted to the distal colon and rectum in AOM-DSS, whereas they occur more proximal in the *APC*-driven model (**Extended Data Fig. 3g**), which may have a greater impact on colonisation.

**Fig. 2.**
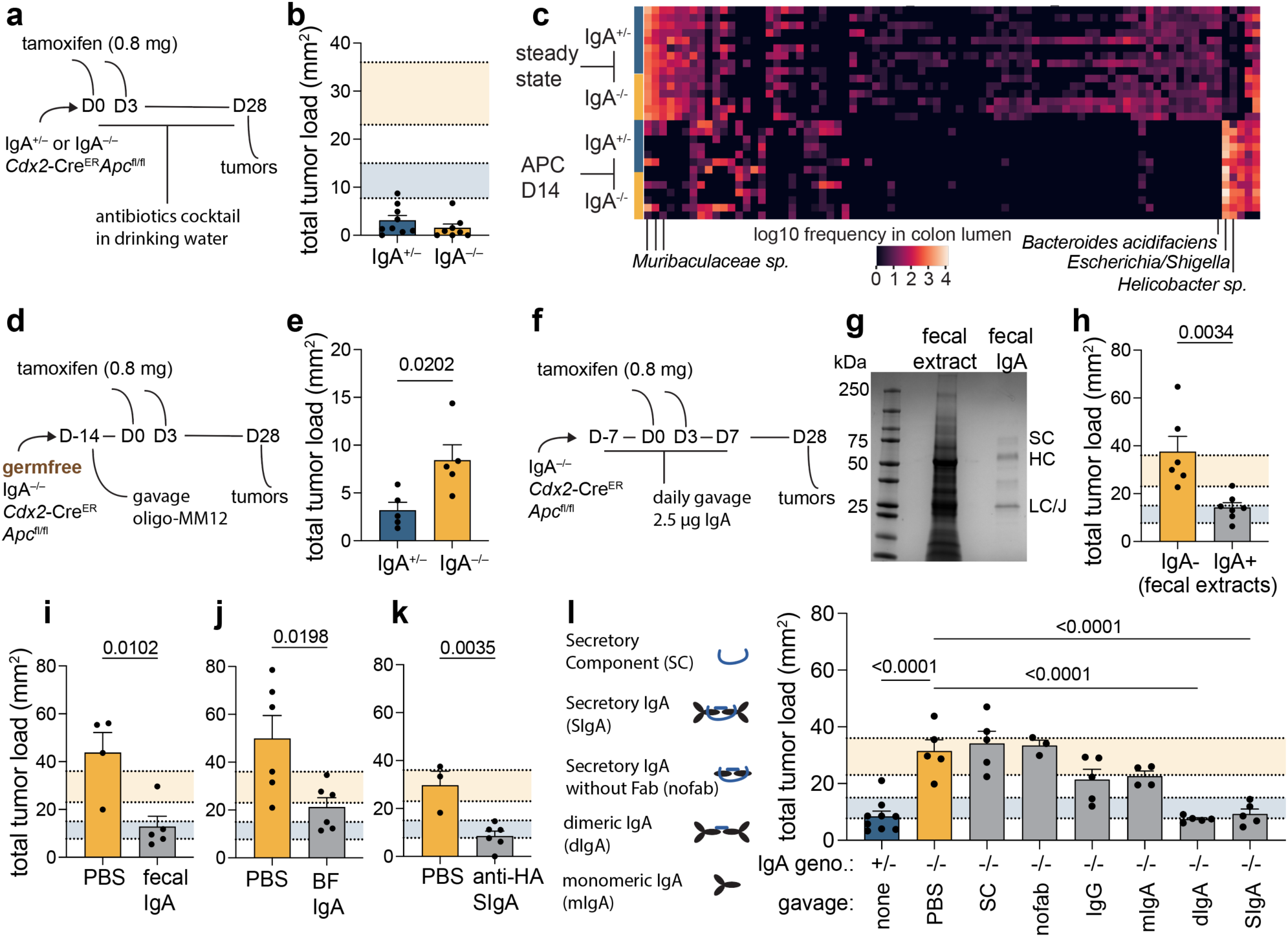
Role of the microbiome and IgA structure in tumourigenesis. **a**, Experimental design for broad spectrum antibiotic treatment (ampicillin, vancomycin, neomycin, metronidazole) to assess the role of microbiome in tumour development in the context of IgA deficiency and *APC* loss. **b**, Total tumour load was dramatically reduced by antibiotic treatment in both IgA^+/−^ and IgA^−/−^ mice to similar levels (pooled 2 cohorts, n = 9, 8). **c**, Heatmap of amplicon sequence variants (columns) identified in colon lumen samples from indicated mice (rows) demonstrates dramatic shift caused by *APC* loss, but no effect of IgA (pooled 2 cohorts, n = 9, 6, 7, 6). **d**, Experimental design for gnotobiotic colon cancer model with a defined microbial community (oligo-MM12) and **e**, corresponding tumour load (pooled 2 cohorts, n = 5). **f**, Experimental design for gavage treatments to prevent tumourigenesis in IgA^−/−^ mice in the context of *APC* loss. **g**, Denaturing gel of sterile faecal extract and resulting purified faecal IgA. **h-k**, Total tumour load following *APC* loss in IgA^−/−^ mice treated with **h**, indicated sterile faecal extracts from healthy donor mice (pooled 2 cohorts, n = 6), **i**, purified faecal IgA from healthy donor mice (pooled two cohorts, n = 4, 5), **j**, purified faecal IgA from germfree mice mono-colonized with *Bacteroides fragilis* (pooled two cohorts, n = 6), **k**, recombinant CR9114 anti-influenza HA SIgA (n = 3, 6), or **l**, structural variants of recombinant antibody (n = 9, 5, 5, 3, 5, 4, 5, 5). (Mean and SEM, shaded areas in tumour load plots represent the 95% confidence interval of tumour load for IgA^+/−^, blue, and IgA^−/−^, yellow, mice from Fig. 1, two-tailed t-tests or one-way ANOVA, p values hidden when p > 0.1)

To further test whether the regulation of specific bacterial species is responsible for the effect of IgA on tumour susceptibility, IgA^+/−^ and IgA^−/−^ *Cdx2^CreER^APC^fl/fl^* mice were re-derived in germ-free isolators by c-section and colonised with a twelve species model microbiome prior to inducing *APC* loss (**Fig. 2d**), which was sufficient to induce normal faecal IgA levels (**Extended Data Fig. 4a**). Here, with acute colonisation by a defined microbiome lacking pathobionts such as *Shigella* and *Helicobacter*, IgA^−/−^ littermates still developed more tumours (**Fig. 2e**). Collectively, these experiments, involving both the measurement and manipulation of the microbiome, are consistent with IgA acting on a pathway that is dependent on bacterial colonisation but does not relate to the control of particular pro- or anti-tumour species.

To rule out secondary effects of IgA deficiency, and determine whether antibody specificity is involved, oral treatments were tested for their ability to rescue IgA^−/−^ mice (**Fig. 2f**). First, sterile-filtered faecal extracts (containing proteins and soluble metabolites, **Fig. 2g**) from healthy mice were gavaged daily starting prior to the induction of *APC* loss. Although treatment was ceased 3 weeks prior to measuring tumours, extracts from IgA^+/−^ but not IgA^−/−^ mice were sufficient to reduce the tumour load in IgA^−/−^ recipients (**Fig. 2h**). To test whether this impact was conferred by the antibodies or other components of the extract, secretory IgA was purified from the faeces of healthy mice (**Fig. 2g**) and gavaged daily in the same timeframe. This polyclonal IgA preparation from healthy mouse faeces was also sufficient to protect IgA^−/−^ mice from increased tumours (**Fig. 2i**). We next tested the role of antibody specificity, first by purifying IgA from the faeces of germ-free Swiss Webster mice mono-colonised with *Bacteroides fragilis*. Despite this restricted mucosal antigen exposure, IgA from these mice was sufficient to protect IgA^−/−^ mice from increased tumour load (**Fig. 2j**). Analyses of recipient faeces during this treatment revealed no coating of the microbiota (**Extended Data Fig. 5a**), raising the possibility that the effect on tumourigenesis is independent of bacterial binding. Consistent with this, recombinant monoclonal secretory IgA specific for influenza HA (CR9114) also protected IgA^−/−^ mice from increased tumour formation (**Fig. 2k**). The functionality of this recombinant antibody allowed us to investigate the structural requirements for tumourigenesis suppression. While either secretory or dimeric IgA treatment was sufficient to provide protection, neither the secretory component nor the truncated secretory IgA without Fabs was sufficient (**Fig. 2l**). Monomeric IgA and IgG treatments did not result in a statistically significant impact on tumour load (**Fig. 2I**). Oral treatments that reduced tumour load mainly reduced the number rather than the size of tumours (**Extended Data Fig. 5b-e**). The dependence of this impact on antibody structure, but not antigen specificity, led us to investigate the potential impact of IgA directly on the endogenous population of cells that become tumours: the colonic epithelium.

## Impact of IgA on epithelial cells

To investigate how IgA may affect the steady-state epithelium and influence the probability of tumourigenesis, mid-colon epithelial cells were sorted (live CD45- EPCAM+, **Extended Data Fig. 6a**) for single-cell transcriptomics from three IgA^+/−^ mice and three IgA^−/−^ littermates (**Fig. 3a-b**). Clusters were identified based on known markers, such as stem (*Lgr5*+) and transit-amplifying (*Mki67*) cells partially overlapping in clusters 0, 5, and 9, and goblet cells in clusters 6, 8, and 10 expressing the secretory progenitor marker *Atoh1* (**Fig 3a-b**, **Extended Data Fig. 6b**). In terms of cellular composition, IgA^−/−^ mice had more cells in clusters 1 and 5 (**Fig. 3c-d**). Cluster 5 represents a population of cycling or early post-mitotic progenitors and transit-amplifying cells, expressing cycling markers *Mki67* and *Nox1* and the canonical Notch target and absorptive lineage identifier *Hes1*^43^ (**Fig. 3b**). Lack of *Muc2* expression further indicated their undifferentiated state (**Extended Data Fig. 6b**). Cluster 1 represents a mature population of colonocytes, though they lack expression of *Aqp8*, the canonical marker for absorptive cells. Instead, these cells, as well as their intermediate cluster 4, were marked by the expression of antimicrobials *Dmbt1*, *Reg3b*, and *Reg3g* (**Fig. 3b, d**), indicating an immunological role. In terms of gene expression, while only a handful of genes were differentially expressed when analysing all cell types together (**Fig. 3e**, **Supplementary Data 3**), these three antimicrobials were among the genes upregulated in colonocytes from IgA^−/−^ mice (**Fig. 3e, Extended Data Fig. 6c, e**). These specific genes were previously found to be epithelium-induced at the single-cell level by bacterial colonisation of germ-free mice^44^. Considering pathways, epithelial cells from IgA^−/−^ mice showed a decrease in interferon signalling and an increase in antimicrobial responses, but not a general inflammatory response (**Fig. 3f, Extended Data Fig. 6d**). Pathways related to proliferation and central metabolism were upregulated in IgA^−/−^ mice, whereas adhesion and tight junction pathways showed cell type-specific changes (**Fig. 3g**). Cluster 9 showed enrichment of the pre-Notch reactome in IgA^−/−^ mice, consistent with Notch induction accompanying the shift of these cycling cells toward the absorptive fate of the expanded cluster 5 (**Fig. 3g**). Collectively, the analysis of the colon epithelial transcriptome at the single-cell level in IgA^−/−^ mice revealed specific changes in epithelial development related to the expansion of the absorptive progenitor population and enhanced antimicrobial function.

**Fig. 3.**
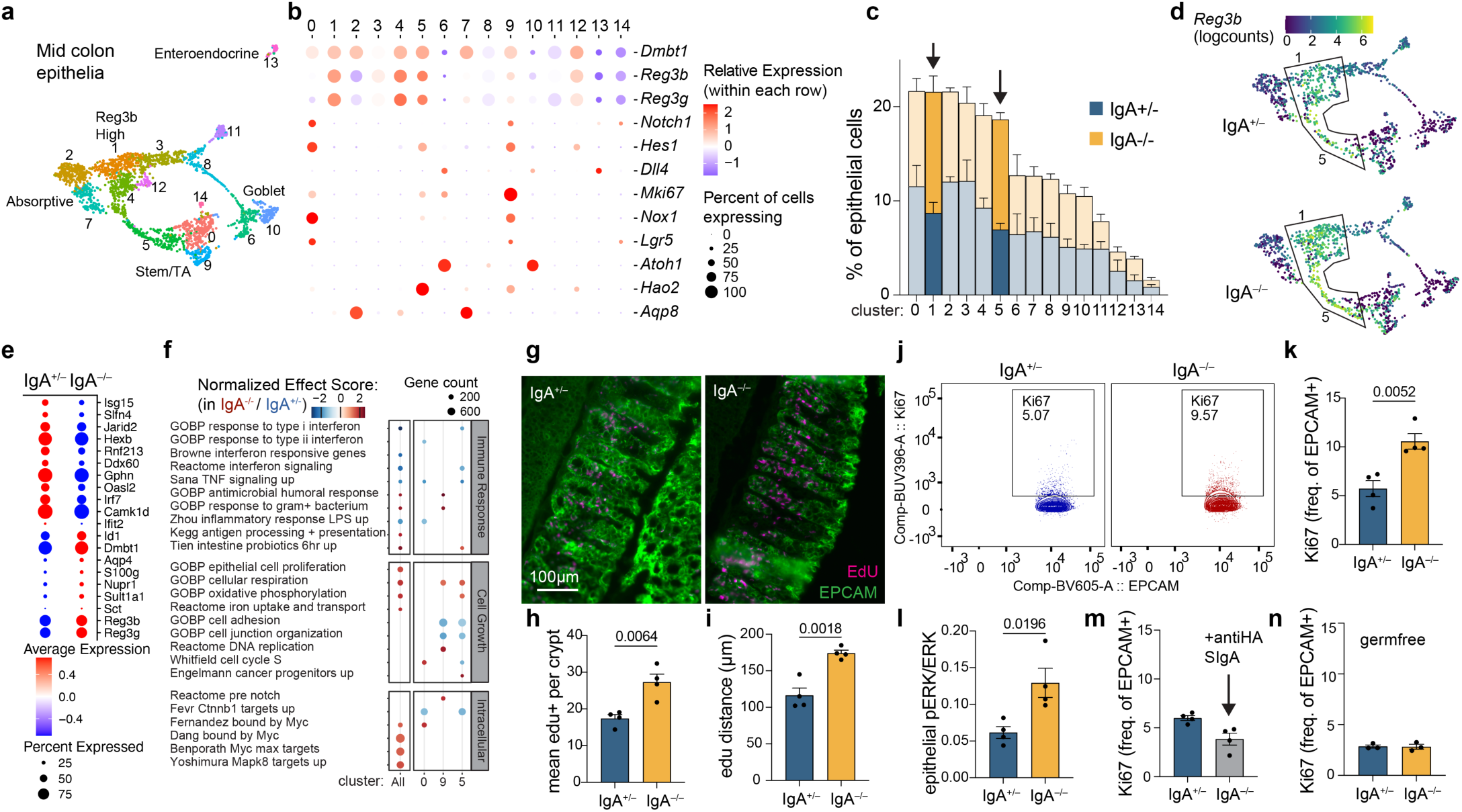
Cellular changes in the colon epithelium of IgA^−/−^ mice. **a**, UMAP and clustering of sorted single colon epithelial cell (live, CD45-, EPCAM+) transcriptomes from IgA^+/−^ and IgA^−/−^ littermates (n = 3). **b**, Enrichment of genes of interest across clusters. **c**, Frequency of each cluster of cells, revealing increases in cluster 1 and 5 in IgA^−/−^ mice (n = 3). **d**, Expression of *Reg3b*, the most differentially expressed gene between IgA^+/−^ and IgA^−/−^ littermates is also associated with cluster 1 and 5. **e**, Top differentially expressed genes depending on IgA genotype across all colonic epithelial cells. **f**, Gene set enrichment analyses across all cells and within individual clusters based on IgA genotype revealed a switch from interferon signalling to anti-bacterial pathways, increases in cell cycling, and alterations in intracellular signalling processes. Dots are visible where adjusted p value < 0.05, size of dot represents the number of genes in the pathway, and color represents the normalized enrichment score of that pathway (IgA^−/−^-enriched is positive, red). **g**, Fluorescence microscopy of frozen sections of swiss-rolled mid-colon from mice injected with EdU at 24 and 4 hours prior. **h**, Quantification of EdU positive cells per crypt, and **i**, maximum distance of EdU positive cells from the crypt base, averaged across at least 20 crypts and 5 frames of view per animal (n = 4). **j**, Flow cytometry of colon epithelial cells (CD45-, EPCAM+) with intracellular staining of the cell cycle marker Ki67. **k**, Quantification of the frequency of Ki67+ epithelial cells depending on IgA genotype (n = 4). **l**, ELISA of lysate of isolated colon epithelial cells specific for phosphorylated ERK normalized to the level of total ERK (n = 4). **m**, Ki67 assessment by flow cytometry as before, but IgA^−/−^ mice were gavaged daily with 2.5 µg SIgA for one week (n = 3, 4), or **n**, were re-derived germfree (n = 3) (mean and SEM, two-tailed t-tests, p values hidden when p > 0.1).

We directly tested whether the developmental and cellularity changes observed by single-cell transcriptomics could be observed by measuring epithelial proliferation. Mice were injected with 5- ethynyl-2-deoxyuridine (EdU), which is incorporated into the DNA of dividing cells, 24 and 4 hours before tissue harvest to label all proliferating cells. Using a counterstain for EPCAM (green) to visualise crypt structures, EdU+ cells (pink) were quantified in the frozen sections (**Fig. 3g**). Sections from IgA^−/−^ mice had increased EdU+ cells per crypt (**Fig. 3h**) and a greater distance travelled from the crypt base (**Fig. 3i**), both consistent with increased epithelial proliferation. Isolated epithelial cells from IgA^−/−^ mice also had a higher frequency of intracellular Ki67 measured by flow cytometry (**Fig. 3j, k**), and a higher ratio of phosphorylated ERK as measured by ELISA (**Fig. 3l**), consistent with accelerated epithelial proliferation. Gavage treatment of IgA^−/−^ mice with monoclonal anti-influenza SIgA was sufficient to reduce Ki67 staining to the level observed in IgA^+/−^ littermates (**Fig. 3m**). As colon epithelial cycling was previously shown to be a physiological response to microbial stimulation through TLR signalling^45^, germfree IgA^−/−^ mice were used to test whether enhanced epithelial proliferation was dependent on the microbiota. In these animals, no IgA-dependent differences were observed in Ki67+ cells measured by flow cytometry (**Fig. 3n**) or EdU+ cells imaged in frozen sections (**Extended Data Fig. 7a**). This set of experiments establishes that IgA controls colon epithelial cell cycling in mice through a pathway dependent on the microbiota but not antibody specificity.

Though not precedented, IgA may plausibly impact epithelial proliferation through a cytokine-like function, rendering antigen specificity irrelevant. To test this, colonic crypts were isolated from IgA^−/−^ mice for organoids. Subsequent treatment with SIgA, but not IgG, reduced the number of recovered organoids (**Extended Data Fig. 7b**). Growth of a mouse colon epithelial cell line (CMT-93) was slowed by treatment with SIgA but not IgG, monomeric IgA, secretory component, or no-fab SIgA (**Fig. 4a**). Several putative immunoglobulin receptors have been reported on colon epithelial cells, but none are known to bind the secretory form of IgA or regulate proliferation. To obtain an unbiased view of the scope of secretory IgA interacting partners expressed by these cells, mouse colon organoid lysates were probed with SIgA, which was then purified using magnetic beads. Co-precipitated proteins were analysed using mass spectrometry (**Fig. 4b**). Compared to controls without SIgA, 54 proteins were significantly enriched (**Supplementary Data 4**), including the primary components of SIgA: the alpha heavy chain, kappa light chain, secretory component, and jchain. Of the 28 non-immunoglobulin-related enriched proteins, most were intracellular, but two surface proteins were highly enriched: scavenger receptor DMBT1 and glycoprotein receptor GP2. Interestingly, *Dmbt1* was one of the three most highly upregulated genes in the colon epithelial cells of IgA^−/−^ mice (**Fig. 3e**), and its expression was highest in the absorptive progenitor and antimicrobial colonocyte populations that were expanded in IgA^−/−^ mice (*see* **Fig. 3b, c**). To test whether these candidate immunoglobulin-binding surface proteins are responsible for SIgA-dependent growth suppression, guide RNAs were designed for each in a targeted CRISPR screen that included *Pigr*, *Dmbt1*, and *Gp2* (**Fig. 4c**). As expected, most of the generated CMT93 cells exhibited slower growth when incubated with SIgA compared to IgG (**Fig. 4d**), but two guide RNAs eliminated this phenotype, both of which targeted *Dmbt1*.

**Fig. 4.**
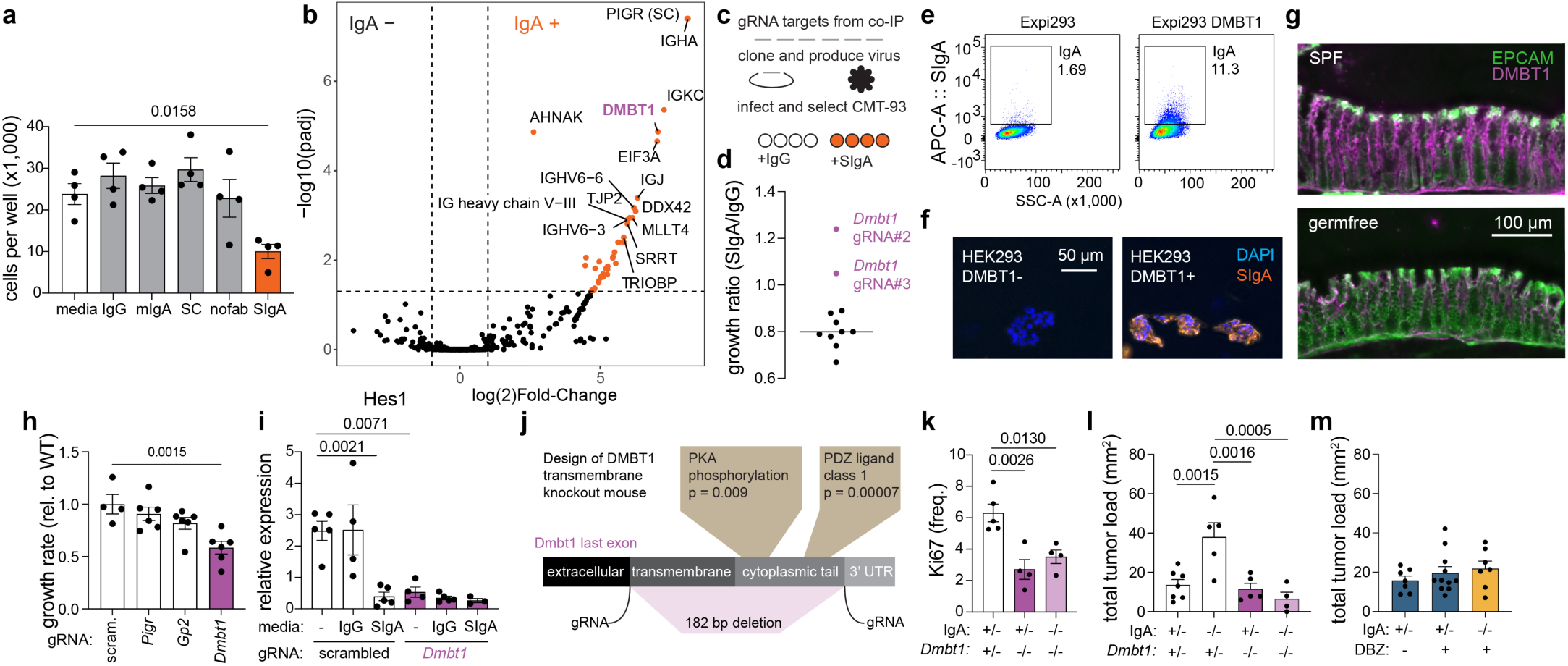
SIgA suppresses epithelial proliferation and tumourigenesis via DMBT1. **a**, CMT-93 mouse colon epithelial cell line cultured with recombinant antibodies for 5 days (n = 4, representative of two independent experiments). **b**, Purified secretory IgA pulldown and mass spectrometry with epithelial organoid lysate (n = 3). **c-d**, Targeted CRISPR screen to discover epithelial surface proteins that are required for regulation of proliferation identifies DMBT1. **e**, Exogenous expression of *Dmbt1* in Expi293 cells confers the ability to bind secretory IgA on the cell surface as assessed by flow cytometry in cells grown in suspension or **f**, imaging of adherent HEK293 cells (representative of 3 experiments). **g**, Immunofluorescence of DMBT1 in frozen sections of colon (representative of 3 mice). **h**, Growth rate (as a fraction of wildtype) of CMT-93 cells with CRISPR knockouts of indicated genes (n = 4, 6, 6, 6). **i**, qPCR of canonical Notch target, *Hes1*, in CMT-93 cells treated with indicated antibodies for 24 hours (n = 5, 4, 5, 4, 5, 3). **j**, *Dmbt1* last exon genetic structure, intracellular signalling potential, and strategy for a novel mouse knockout of the transmembrane and signalling domains. **k**, Colon epithelial cell proliferation as assessed by flow cytometry of intracellular Ki67 staining in isolated colon epithelial cells from *Dmbt1* signalling knockout mice (pooled two cohorts, n = 5, 4, 4). **l**, Assessment of tumour outcomes four weeks after epithelial-specific loss of *APC* induced in IgA and *Dmbt1* single and double knockouts (pooled two cohorts, n = 7, 5, 5, 4). **m**, total tumour load of mice treated with the Notch inhibitor, dibenzazepine (pooled two cohorts, n = 7, 11, 7)(mean and SEM, one-way ANOVA, p values hidden when p > 0.1).

## IgA regulates epithelia through DMBT1

*Dmbt1* (deleted in malignant brain tumours) encodes several large (∼200 kDa), highly glycosylated mucin-like proteins. The human protein was originally described as a bacterial agglutinin and was co-purified as a complex with IgA in saliva^46^. The gene was independently discovered years later and was found to encode a single transmembrane domain on the final exon in both mice and humans^47^. The protein has been observed on both apical and basolateral sides of mid-crypt colonocytes in humans^48^ and mice^49^, but its functions are enigmatic. To begin to study this, Expi293 cells were engineered to overexpress the transmembrane protein. Surface expression was confirmed by flow cytometry (**Extended Data Fig. 7c-d**) and was sufficient to confer cell surface binding of SIgA (**Fig. 4e**). This was reproduced independently in adherent HEK293 cells grown on a coverslip and assessed for their capability to bind SIgA using imaging (**Fig. 4f**). DMBT1 is not a specific receptor for SIgA, as it also binds IgG and dimeric IgA, as shown by flow cytometry (**Extended Data Fig. 7d-e**), consistent with its previously shown binding promiscuity^50^. Microscale electrophoresis of purified DMBT1 protein confirmed relatively low affinity (hundreds of nM to low µM Kd) interactions with SIgA, dimeric IgA, monomeric IgA, and IgG (**Extended Data Table 1**). Notably, in the colon lumen, no isotype reaches the high concentration of SIgA (**Extended Data Fig. 1c-e**), so functional binding to IgA *in vivo* may relate more to accessibility rather than biochemical affinity. In mice, microbiota-dependent DMBT1 protein was evident in the colon epithelium by immunofluorescence staining (**Fig. 4g**), although staining was ubiquitous in tissue, likely due to secreted forms of the protein.

Next, we explored the relationship between DMBT1 and epithelial proliferation. *Dmbt1*^−/−^ epithelial cell lines showed slowed growth (**Fig. 4h**), consistent with previous studies in which *Dmbt1* expression was positively associated with proliferation in normal colon^51^ and lung epithelia^52^. As our analysis of IgA impacts on colon epithelial cells implicated the growth of absorptive progenitors that are known to express Notch target genes like *Hes1*^53^ (**Fig. 3b-f**), we assessed *Hes1* expression as a molecular correlate of proliferation. *Hes1* expression was substantially decreased upon *Dmbt1* knockout (**Fig. 4i**). Additionally, treatment of wild-type cells with SIgA, but not IgG, decreased *Hes1* expression (**Fig. 4i**). However, treatment of *Dmbt1*^−/−^ cells with SIgA had no further effect, placing DMBT1 downstream of SIgA in a pathway that regulates proliferation.

To specifically interfere with its role as a membrane protein *in vivo*, we designed a mouse carrying a mutation in the last exon of *Dmbt1* (which is excluded from secreted isoforms), removing its transmembrane domain and cytoplasmic tail with a potential signalling function (**Fig. 4j, Extended Data Fig. 8a**). We used two guide RNAs to generate this transmembrane knockout directly in IgA^−/−^ and IgA^−/−^ *Cdx2^CreER^APC^fl/fl^*mice. *Dmbt1* mutation did not affect plasma cells or IgA production (**Extended Data Fig. 8b-g**). However, compared to littermate heterozygous controls, *Dmbt1* ^−/−^ mice showed decreased epithelial proliferation (**Fig. 4k**). Double knockout mice (*Dmbt1* and IgA) showed no increase in proliferation previously observed in IgA deficiency (**Fig. 4k**), indicating that DMBT1 is required for IgA regulation of proliferation. Finally, we tested the role of DMBT1 in IgA^−/−^ mice susceptibility to *APC*-driven tumours. While IgA^−/−^ mice still exhibited increased tumours in a *Dmbt1*^+/−^ context, knockout of transmembrane *Dmbt1* completely reverted IgA^−/−^ susceptibility (**Fig. 4l**). This supports a model whereby IgA binds to DMBT1 and suppresses a previously undescribed mitogenic or pro-proliferative function of this scavenger receptor, consequently reducing tumour chance.

This model predicts that the susceptibility of IgA^−/−^ mice to colorectal tumours is dependent on increased epithelial proliferation during homeostasis. Although *Hes1* expression correlated with SIgA regulation of epithelial growth, knocking out *Hes1 in vitro* was not sufficient to eliminate the effects of SIgA on growth (**Extended Data Fig. 7f**). Still, if increased proliferation in IgA^−/−^ animals drives their tumour risk, inhibiting the pro-proliferative pathway further upstream of IgA-DMBT1 interaction should be protective: gamma-secretase inhibitors such as dibenzazepine (DBZ) are known to block Notch receptor cleavage and reduce proliferation of intestinal epithelial cells. We treated IgA^+/−^ or IgA^−/−^ *Cdx2^CreER^APC^fl/fl^* mice with a sub-therapeutic dose (3 µmol / kg) previously shown to have no impact on tumour growth and minimal impact on normal colon crypts^54^ (**Extended Data Fig. 7g**). While this low-dose treatment 5 days before and 5 days after APC loss did not affect the tumour outcome in IgA^+/−^ mice, their IgA^−/−^ littermates were now indistinguishable in tumour outcome (**Fig. 4r**). This suggests that the outgrowth of Notch-dependent absorptive progenitors during steady state, initially observed by single-cell RNA-Seq (in **Fig. 3**), is responsible for increased tumourigenesis in IgA^−/−^ mice.

In comprehensive published datasets spanning body tissues, *Dmbt1* expression was remarkably restricted to subsets of epithelial lineages in both mice^55^ and humans^56^ (**Extended Data Fig. 10a-b**). Flow cytometry confirmed the presence of DMBT1 on the surface of colon epithelial cells (**Fig. 5a-b**), which strongly correlated with the proliferation marker Ki67 (**Fig. 5c**). Colon epithelial cells from germ-free mice exhibited less Ki67(**Extended Data Fig. 9c**), consistent with previous work^45^, and we found that this was associated with decreased DMBT1 expression (**Fig. 5a-b**). This is consistent with the notion that IgA controls proliferation via a microbiota-induced pathway, potentially explaining why IgA^−/−^ mice display no differences in colon epithelial proliferation under germ-free conditions (**Fig. 3n**). In SPF IgA^−/−^ mice, where enhanced proliferation was observed, this was associated with an increase in DMBT1 expression (**Extended Data Fig. 9d-f**). Additionally, the association of DMBT1 with proliferation was lost in cells isolated from tumour-containing colon tissues from the *APC*-driven cancer model (**Fig. 5a-c**), which showed an overall decrease in DMBT1 (**Fig. 5b**) and an expansion of the DMBT1-negative Ki67+ population (**Fig. 5c**). Based on this, we hypothesized that loss of DMBT1 might specifically mark tumour cells in this model, as suggested in a recent proteomic study of human colorectal polyp biopsies^57^.

**Fig. 5.**
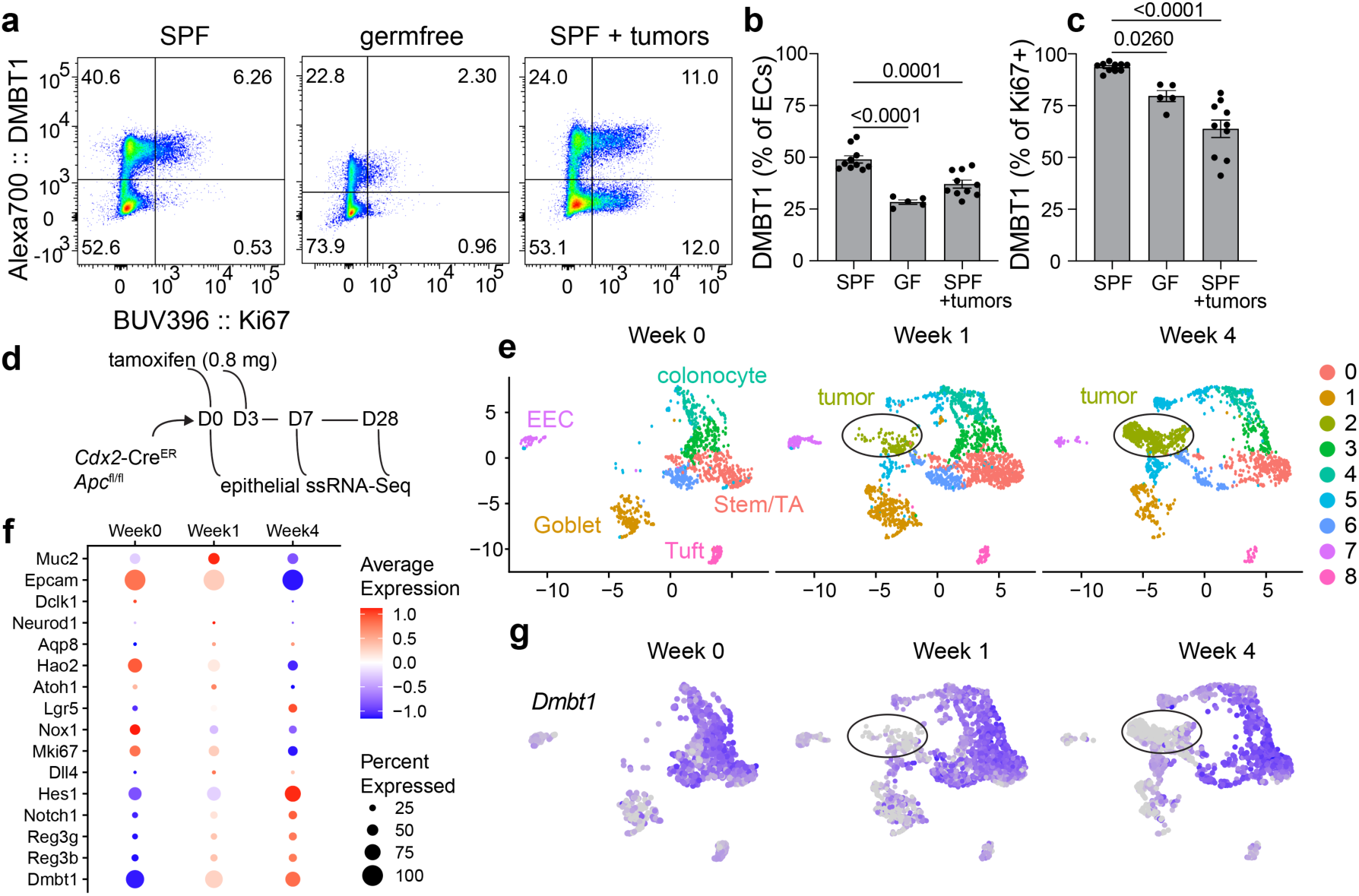
DMBT1 expression is associated with physiological epithelial proliferation and lost during tumourigenesis. **a**, Flow cytometry measurement of surface DMBT1 and intracellular Ki67 in epithelial cells (live, CD45-EPCAM+) from SPF and germfree mouse colons, and tumour-containing colon tissues isolated 4 weeks after *APC* loss, summarized as the DMBT1+ frequency out of **b**, all epithelial cells, or **c**, proliferating epithelial cells (n = 10, 5, 10, mean and SEM plotted, one-way ANOVA, p values hidden when p > 0.1). **d**, Experimental schematic for timecourse single-cell RNA-Seq of colon epithelial cells during APC-driven tumourigenesis. **e**, UMAP of all live, CD45-, EPCAM+ cells sequenced split by timepoint, where cluster 2 marks tumour cells. **f,** Differential expression over time, for markers of colonocyte cell types show overall loss of epithelial identities, increase in stemness, and increase in antimicrobials (*Reg3b*, *Dmbt1*). **g,** UMAP showing loss of *Dmbt1* expression specific to tumour cells. (mean and SEM, one-way ANOVA, p values hidden when p > 0.1).

To take advantage of the temporal control enabled by animal models, we performed a time-course single-cell RNA-seq of the region of the transverse colon where tumours typically emerge (**Fig. 5d**). A tumour population in cluster 2, absent at week 0, was discernible just one week after *APC* loss (**Fig. 5e, Extended Data Fig. 9g**). These cells acquired stemness, with higher *Lgr5* expression than actual stem cells (**Extended Data Fig. 9h**). Considering gene expression changes over time across all populations, epithelial identities were lost, whereas the previously identified antimicrobial module (*Dmbt1*, *Reg3b*, and *Reg3g*) was induced (**Fig. 5f**). Interestingly, colorectal tumour development coincided with an overall increase in *Dmbt1* expression (**Fig. 5f**), but the increase was restricted to untransformed cells, whereas the tumour cells themselves showed a dramatic loss of *Dmbt1* (**Fig. 5g, Extended Data Fig. 9h**). We conclude that *Dmbt1* expression marks physiological epithelial proliferation of the absorptive progenitor population. Ordinarily, this subjects cells to regulation by SIgA; however, tumours may escape this mechanism.

## Discussion

We found that by modulating DMBT1-induced proliferation of colon epithelial cells, IgA in the gut lumen lowers the risk of colon tumour initiation in mouse models. DMBT1 is itself an enigmatic protein (see **Supplementary Discussion**), previously implicated in cancer through undescribed mechanisms.

While microbial stimulation of epithelial cells provides an “accelerate” signal for proliferation and antimicrobial responses, IgA feedback through DMBT1 appears to be a “brake,” closing a homeostatic loop of host responses to the microbiota. A recent study independently reported the susceptibility of IgA^−/−^ mice to AOM-DSS-induced tumours^58^. IgA was previously reported to suppress ovarian tumour growth through interactions with immune cells and pIgR, which is highly expressed in ovarian tumours^59^. In contrast, pIgR is dramatically downregulated in human colorectal tumours^60^, and we observed an effect of IgA only on physiological cell growth (not on tumour growth) with no dependence on *Pigr*. IgA has many different functions which can be tissue and context-specific, and our findings do not supersede other studies thereof. The impacts of IgA on microbiota, as well as on epithelia described here, may intersect with other genetic and environmental factors to manifest in the variable outcomes observed in IgA deficiency^4,7,10,11^.

## Methods

### Mice

All mice were housed at the Rockefeller University or Yale University and handled in accordance with protocols approved by the Institutional Animal Care and Use Committee (IACUC) and under the ethical guidelines within the National Institutes of Health Guide for the Care and Use of Laboratory Animals. C57BL/6J (000664) and *Cdx2^CreER^* (022390) mice were purchased from Jackson Laboratories. *IgA*^−/−^ mice (1999 version) on a C57BL/6J background^33^ were provided by Margaret Conner (Baylor). *IgA*^−/−^ mice (2026 version) on a BALB/c background^40^ were provided by Hans Oettgen (Harvard). *APC^fl/fl^* mice on a C57BL/6J background were provided by Sergei Grivennikov (Cedars-Sinai). *Aid^Cre^* mice^61^ were provided by C-A Reynaud and J-C Weill (Université Paris-Descartes). Mice were housed in Specific Pathogen Free conditions on a fixed 12 hour light-dark cycle, temperature 68-79F, humidity 30-70%, with food and water provided *ad libitum*. C57BL/6J *IgA*^−/−^ mice were re-derived by c-section into germfree flexible film isolators (Class Biologically Clean) that were routinely monitored by quantitative real time PCR. Germfree mice were housed with a fixed 12 hour light-dark cycle with autoclaved food and water provided *ad libitum*. The genotype of mice was determined by quantitative real time PCR analyses by Transnetyx. The IgA genotype of mice was routinely cross-checked using ELISA or flow cytometry. For experiments comparing IgA^+/−^ to IgA^−/−^, littermate, sex-matched controls were co-housed from weaning through the duration to normalize microbiome^62^. Sample sizes were chosen based on pilot data. During data collection, mice were identified only by an experiment name, cage number, and ear punch, which was matched to genotype post hoc. Experiments were initiated when mice were aged 8-10 weeks, unless otherwise noted. Both male and female mice were used for all experiments, analyzed separately, and then pooled if no sex differences emerged. Estrous cycle was not controlled for.

### Generation of transmembrane domain knockout of *Dmbt1*

The knockout of the transmembrane domain of *Dmbt1* was made at Rockefeller University using the Easi-CRISPR method^63^. Briefly, two guide RNAs (AAGTGAAACAACCCAGACAT and ATCCTCCACTGCCAAGTCTG) spanning the transmembrane and cytoplasmic tail region of the last exon of *Dmbt1* were complexed with Cas9 and injected into pronuclei of fertilized embryos harvested from *IgA*^−/−^ *Cdx2^CreER^ APC^fl/fl^* mice. These embryos were implanted into pseudopregnant mice. After sequence confirmation of the excision of 182 bp, founder animals were backcrossed to the parental strain to regenerate the wildtype allele.

### Murine cancer models

For all cancer models, animals were monitored daily for sickness behavior, body condition, and weighed at least twice per week to assess health and maintain a humane endpoint if weight loss exceeded 20% of their initial weight or tumour size was beyond 10 mm (this size limit never occurred in our models). For the APC model, *Cdx2^CreER^APC^fl/fl^*mice were injected intraperitoneally at day 0 and day 3 with 0.8 mg tamoxifen (Sigma) from a stock solution at 4 mg/ml dissolved in corn oil (Sigma). After 4 weeks (day 28), mice were sacrificed to count and measure tumours. For the AOM-DSS model, mice were injected intraperitoneally with 12.5 mg/kg azoxymethane (AOM) at day 0. Exposure to 1.5% dextran sodium sulfate (DSS) in drinking water replaced their normal water for 1 week starting at day 0. This was followed by a 2 week break, a second DSS exposure for 1 week, a 2 week break, and a third exposure for 1 week. Animals were sacrificed at day 100 to measure and count tumours. Lipocalin-2 was measured using the Mouse Lipocalin-2/NGAL DuoSet ELISA kit from R&D Systems (Cat. DY1857) according to manufacturer’s protocol. For the pre-cancerous organoid implant model, IgA^+/−^ and IgA^−/−^ mice were injected with 1.5 x10^5^ cells of AKPS organoids (*APC*, *Kras*(G12D), *P53*, *Smad4*; generated in Sergei Grivennikov’s laboratory). Cells were injected over 3 injections (5 x10^4^ cells in 40 uL each), spaced ∼ 0.5 cm apart in the distal colon and rectum wall using a colonoscope (Karl Storz, Model TC300) with a custom-made 33-gauge needle (Hamilton Company). AKPS organoids were cultured by resuspending organoid cells in 50% matrigel (Corning) in cold ADF^+^ media (Advanced DMEM F12 (Invitrogen) supplemented with 5% heat inactivated FBS (Sigma-Aldrich), 1% Penicillin-Streptomycin-Glutamine (Invitrogen), 1% HEPES (Invitrogen)). In a 6-well plate, 40 uL droplets of cell suspension were plated and incubated at 37°C for 20 minutes, followed by adding 3 mL of pre-warmed ADF^+^ media. To prepare cells for injections, AKPS organoids were dissociated with cold ADF^+^ media to break down matrigel, and spun down at 350g for 5 minutes. Organoid pellet was resuspended in Tryple Express Enzyme (Thermo Fisher Scientific) and incubated at 37°C for 30 minutes with shaking at 120 rpm. Organoids were further broken down into cells by pipetting multiple times with P1000, followed by washing with ADF^+^ media and spinning down at 350g for 5 minutes. Cells were resuspended in ADF^+^ media for counting, then volume was further adjusted to keep cells at 1250 cells/uL in 10% matrigel, 1% filter-sterilized Methylene Blue (Biopharm, Amazon), and 10 uM Y27632 (Stemcell Technologies). Implanted tumours were monitored every two weeks using the colonoscope and were scored according to a previously published system (1: small but detectable tumour, 2: tumour covering one-eighth of colon circumference, 3: tumour covering one-fourth of colon circumference, 4: tumour covering one-half of colon circumference)^64^.

### Interventions in APC cancer model

For antibiotic depletion of the microbiome, mice were provided with sterile-filtered drinking water with 1 mg/ml ampicillin, 1 mg/ml neomycin, 0.5 mg/ml vancomycin, and 0.5 mg/ml metronidazole, supplemented with 10 mg/ml sucralose for palatability. For oral gavage treatments to rescue IgA deficiency, mice were subjected to a daily intra-gastric gavage with 2.5 µg of IgA (or the equivalent) in 200 µl PBS. For Notch inhibition, mice were given a daily 3 µmol/kg intraperitoneal injection of dibenzazepine (DBZ) (StemCell Technologies) which was first dissolved as a 10 µmol/ml in DMSO stock solution (kept frozen at −20C) and then diluted in 0.5% hydroxypropyl methylcellulose for injection. For colonization with oligo-MM12^65^, germfree mice were gavaged once with cecal content from previously-colonized mice with all 12 species as previously described.

### Bulk RNA-Seq

Working on ice with one animal at a time until samples were homogenized by bead beating to stabilize RNA, mice were sacrificed by cervical dislocation, the colon was dissected, cut open longitudinally, and washed 3 times in PBS to remove lumen contents. An approximately 0.5 cm x 0.5 cm sample of tissue from the mid-colon (3 cm distal from the cecal junction) was collected in a Lysing Matrix D tube (MP Biomedicals) with 1 ml Trizol (Invitrogen) and beat at max speed for 60 seconds (MP Biomedicals FastPrep 24 Bead Beater), twice, with 5 minutes rest on ice between. Total RNA was purified according to the manufacturer’s instructions for Trizol, followed by further purification using a Qiagen RNeasy column with on-column DNase treatment (Qiagen) according to the manufacturer’s instructions. RNA quality was assessed using an Agilent Bioanalyzer and only samples with RIN > 8.0 were used for poly- A selected mRNA-Seq. Libraries were sequenced on an Illumina NextSeq 500 high-throughput flow cell, with a single-end 75 bp read, yielding between 23 and 30 million high quality paired-end reads per sample. STAR aligner 2.7.10a and biomaRt 2.52.0 were used to align reads and annotate genes with the GRCm39 mouse genome and M32 mouse transcriptome. Differential expression analyses was performed using DESEq2 1.36.0 and gene set enrichment analyses using fgsea 1.22.0 with gene sets c2.cgp.v7.5.1.symbols and c5.go.bp.v7.5.1.symbols.

### 16S rRNA amplicon sequencing

Mice were sacrificed by cervical dislocation and the colon was dissected and opened longitudinally on ice using autoclaved tools and aseptic technique. Lumen samples were taken from the mid colon, then the tissue was washed thoroughly 3 times in sterile PBS on ice, dabbed several times on a sterile petri dish to dry, and mucus was gently scraped from the tissue using a plastic cell scraper. All samples were frozen at −80C prior to further processing. DNA extraction was performed using the QIAamp PowerFecal Pro DNA Kit (Qiagen) with bead beating on an MP Biomedicals Fastprep 24, according to manufacturer’s protocols. To avoid contaminating the lower biomass samples, mucus samples were processed separately, with decontamination of benches, racks, pipettes, and microcentrifuge prior to processing. 16S rRNA was amplified and libraries were prepared and pooled according to the Earth Microbiome Project 16S Illumina Amplicon protocol (earthmicrobiomeproject.org). Sequencing was performed with an Illumina MiSeq 150 bp paired-end run. The data were analyzed using the Qiime 2 (v2022.8) package. Amplicon sequence variants (ASVs) were identified and chimeric sequences were removed using Deblur. Phylogenetic analyses were performed using the Qiime 2 classifier trained on the SILVA database (v138) from 515F to 806R to match the amplicon sequencing. Following the Deblur processing, only samples with more than 1000 reads were included and only features with at least 100 reads across all samples that were present in at least 5 different samples were included. Differential abundance was assessed using a dedicated statistical test (ANCOM)^66^.

### Faecal IgA purification

Similar to a previously published method^67^, the small intestine, cecum, and colon were dissected from three healthy animals, and the contents were pooled into a 15 mL conical tube. To the pooled contents, 6 mL of 1× PBS-T (PBS with 1% Tween 20, prepared from 10× stock (AlfaAesar)) containing a protease inhibitor cocktail (Roche) was added. The mixture was vortexed at maximum speed for 5 minutes. After vortexing, the suspension was centrifuged at 3000 g for 10 minutes. The supernatant was then divided into six 2 mL tubes, which were centrifuged again at 10,000 g for 10 minutes. During this second centrifugation, Protein L magnetic beads (Pierce™ Protein L Magnetic Beads, Thermo Scientific) were prepared. The bottle of beads was gently vortexed to mix, and 200 µL of Protein L beads was added to each of the six 2 mL tubes, followed by the addition of 300 µL of PBS-T. The tubes were vortexed gently for 1 minute, after which the beads were collected using a magnet, and the supernatant was removed. This wash step was repeated by adding 1 mL of PBS-T, vortexing gently for another minute, and collecting the beads with a magnet to remove the supernatant. Following the beads’ preparation, the IgA supernatant was transferred into the tubes with the washed Protein L beads. The mixture was shaken gently at room temperature for 1 hour to allow for binding. After incubation, the beads were collected using a magnet rack, and the supernatant was discarded. The beads were then washed three times with 1 mL of PBS-T, gently vortexed for 1 minute during each wash, followed by collection of the beads with a magnet and removal of the supernatant. To elute the bound IgA, 200 µL of elution buffer (0.1 M glycine at pH 2-2.5) was added to the beads. The tubes were gently vortexed and incubated at room temperature for 10 minutes, with occasional gentle vortexing. The beads were then collected with a magnet, and the supernatant containing the eluted IgA was combined into a new tube. To neutralize the pH of the eluted IgA, an appropriate volume of neutralization buffer (10% 1 M Tris-HCl at pH 8) was added. Concentration was checked using a Nanodrop (for crude measurement) or an ELISA assay. Finally, purified IgA was sterile filtered through a 0.22 µm filter.

### Monoclonal IgA and structural variant generation

Recombinant secretory component, CR9114 SIgA, and all structural variants were produced following previously published methods^68^. Briefly, genes encoding the *mus musculus* IgA heavy chain constant region (Uniprot P01878) and the lambda light chain constant region (Uniprot A0A0G2JE99) were fused with the CR9114 VH and VL domain sequences^69^ to create complete heavy chain and light chain sequences. Alternatively, the *mus musculus* IgA heavy chain constant region domains CH3 and CH4 were fused to an N-terminal hexahistine tag to create a Fc(no fabs) expression construct. The TPA signal sequence (residues MDAMKRGLCCVLLLCGAVFVSPAGA) was encoded at the start of the heavy chain sequences and the mouse IgKappa signal sequence (residues METDTLLLWVLLLWVPGSTG) was encoded at the start of the light chain sequence. These sequences, along with *mus musculus* JC (Uniprot P01592; native signal peptide) and *mus musculus* pIgR ectodomain (SC) residues 1-567 (Uniprot O70570; native signal peptide) with C-terminal hexahistine affinity tag sequence, were cloned into mammalian expression vector pD2610v1 (Atum). Resulting expression constructs were transiently transfected or co-transfected into HEK Expi-293-F cells with the ExpiFectamine 293 transfection kit (ThermoFisher, A14525), according to company protocol. Resulting proteins were purified via CaptureSelect LC-lambda (mouse) affinity matrix (ThermoFisher, 194323010) or Ni-NTA agarose (Qiagen, 30210), and by Superose 6 Increase 10/300 GL (Cytiva Life Sciences, 29091596) size exclusion chromatography using an ÄKTA pure 25 chromatography system (Cytiva Life Sciences). Purified proteins were exchanged into 1x phosphate buffered saline buffer (HyClone, SH30258.01) prior to use in experiments.

### Colon epithelial cell isolation

Mice were sacrificed by cervical dislocation. The colon was dissected on ice and opened longitudinally. Lumen content was gently removed and the tissue was washed 3 times in ice cold PBS. The mid colon was taken for further processing (a length of the tissue starting from 3 cm distal from the cecal junction and continuing for 4 cm toward the rectum). The tissue was cut into 1 cm pieces into PBS with 30 mM EDTA and 1.5 mM DTT and incubated on ice for 20 minutes with a brief shake at the end to remove mucus. The tissue was moved to DMEM with 30 mM EDTA and 2% fetal calf serum and incubated at 37C in a waterbath for 10 minutes with an aggressive shake for 30 seconds at the end to remove epithelial cells from the tissue. The cells in media were collected after passage through a metal sieve to remove remaining tissue. Pelleted cells were washed once with HBSS with 10% FCS then resuspended in HBSS (with divalent cations) with 200 µg/ml dispase II (Sigma D4693) and incubated for 10 minutes at 37C in a water bath with brief shaking every 2 minutes to dissociate epithelia into a single cell solution (confirmed by light microscopy). The reaction was stopped by adding FCS to 10% and mixing by inversion. Cells were filtered through a 70 µm filter and washed with HBSS 10% FCS. Cells were stained for flow cytometry in a complete T cell media: RPMI-1640 (Gibco 21870) plus 10% fetal bovine serum (Gibco10437), 1% Pen/Strep (Gibco15140), 1% L-glutamine (Gibco25030), 1% NaPyruvate (Gibco113690), 1% Nonessential amino acids (Gibco 11140), 1% HEPES (Gibco 15630) and 2- Mercaptoethanol [50uM] (Aldrich M6250). For intracellular staining, cells were fixed, permeabilized, and stained using the eBioscience Foxp3 Transcription Factor Staining Buffer set according to the manufacturer’s recommendations.

### Single cell RNA-Seq

Epithelial cells isolated as described above (Colon epithelial cell isolation) were stained with individual Biolegend TotalSeq C antibodies (1:100) to allow for sample-level barcoding. Epithelial cells (SSC, FSC, live, CD45-, EPCAM+) were sorted on a BD FACSAria III Cell Sorter using a 70 µm nozzle at 45 psi, pooling 20,000 cells per sample together in one tube (total 120,000 cells). Post-sort viability was estimated to be 90% by trypan blue staining. A sample of the cell mixture was loaded into a single well of the 10X Genomics Chromium microfluidic device. A single-cell gene expression library was generated following the manufacturer’s protocol for a Chromium Next GEM Single Cell 5’ Reagent Kit v2 (Dual Index) with Feature Barcode technology for Cell Surface Protein (to capture the TotalSeq C sample barcodes). Libraries were sequenced on a NovaSeq S1 flow cell. Primary analysis, encompassing alignment and generation of feature barcodes matrices, was performed on Cell Ranger (v7.0.1), which showed 2,764 cells captured with a mean of 152,000 reads and 2,915 genes per cell. Cell Ranger was run with the option ‘--feature-ref=’ to add the sequence information of the first six barcodes on TotalSeq C, the first three corresponding to IgA^+/−^ samples and the last three to IgA−/− mice, according to the manufacturer recommendations. Filtered feature barcode matrices were then imported to the R environment using Seurat (v4.3.0) and all further analyses were done in R. Cells were filtered by singlet hashtag status, percentual of mitochondrial gene expression less than 10% and maximum RNA count per cell to avoid doublets and dying cells, then were scaled by SCT transform and all the PCA, UMAP and clustering possibilities were calculated. Final clustering resolution was decided according to biological difference between epithelial cell types and cell type annotation was performed using enrichment packages as ClusterProfiler (v4.6.2) and fgsea (v1.24.0) and manual curation. Differential Expression was perfomed using the FindMarkers function in Seurat. Graphs were plotted using Seurat functions, iSEE visualization package (v2.10.0) and ggplot2 (v3.4.3). Final functional enrichment for figures was done using fgsea (v1.24.0) against GO BP and msigdb mice c2 databases (c2.cgp.v7.5.1.symbols and c5.go.bp.v7.5.1.symbols).

### Measurements of *in vivo* epithelial proliferation

Mice received 500 µg of 5-ethynyl 2’-deoxyuridine (EdU) (Thermo Fisher A10044) in a 100 µl PBS intraperitoneal injection at 24 hours and 4 hours prior to sacrifice. The colon was dissected on ice, opened longitudinally, and lumen content was gently removed with forceps. Swiss rolls of colon tissue were fixed in PBS-buffered 2% PFA for 2 hours at 4C, washed three times with PBS, and frozen in OCT. A cryostat was used to cut 16 µm sections which were adhered to Superfrost Plus White slides (Fisherbrand). Edu was visualized using the Click-iT EdU Alexa Fluor 647 Imaging Kit (Thermo Fisher) according to the manufacturer’s protocol, including a counterstain with 1:200 AF488 anti-mouse CD326 (EPCAM) (Biolegend Cat#118210). Stained sections were mounted with Fluoromount G (Southern Biotech) and imaged on a Keyence BZ-X800 epifluorescence microscope. Crypts were assessed in the mid-colon region (only the middle third of the length of the colon) and only counted when both the bottom and top of the same crypt was clearly discernable by EPCAM staining. At least 5 fields of view and at least 20 crypts were counted for each sample. For Ki67 staining, epithelial cells were prepared from mid colon samples as described above (Colon epithelial cell isolation) and analyzed on a BD FACSymphony. Phosphorylated ERK was measured by preparing epithelial cells as described above (Colon epithelial cell isolation) and using a Phospho-Erk1 (pThr202 / pTyr204 + Erk2 (pTyr185/187) and pan-Erk1 / 2 ELISA Kit from Sigma (Cat. RAB0349) according to manufacturer’s instructions.

### Isolation and culture of mouse-derived colon organoids

Crypts were isolated from the colons of IgA knockout mice through successive 30-minute and 1-hour incubations at 4°C in PBS containing 10 mM EDTA, according to a published method^70^. After isolation, the cells were cultured in a hybrid medium composed of 50% primary medium (DMEM/F12 supplemented with 1% HEPES (Gibco 15630), 1% L-glutamine (Gibco25030), 1% Pen/Strep (Gibco15140), 20% fetal bovine serum (Gibco10437)) and 50% conditioned L-WRN medium, further enriched with 50 ng/mL EGF (Sigma SPR 3196-500). Crypts in hybrid media were mixed with matrigel (Fisher Scientific CB40230C) (30-50%) and dispensed into a 48-well plate, with each well receiving 200 μL of the mixture. The plate was incubated for 20 minutes at 37°C. After this period, 300 μL of enriched hybrid media was added to the top of each well. Following three days, 200 uL of enriched hybrid media was added to each well, and slow growth of cells was observed for two more days. For passaging, organoids were dissociated by incubation in TrypLE™ (Thermo Fisher Scientific 12605010). For subsequent culture, the cells were placed in the enriched hybrid media mixed with matrigel (30- 50%). A volume of 40 μL of the cell-matrigel suspension was dispensed into 5 distributed drops per well in a 6-well plate. The plate was incubated to polymerize the matrigel at 37°C for 5 minutes and then inverted for 15 minutes to prevent cell attachment to the bottom of the plate surface. Treatments with antibodies [10ug/mL], including IgA purified from faeces and mouse IgG2a isotype control (BioXCell BE0085), were embedded in supplemented hybrid media and placed in their respective wells. The media was changed after 3 days. Imaging was performed for 4 days after adding treatments using Z-stack on a Keyence BZ-X800 epifluorescence microscope. Image quantifications were made manually using ImageJ 2.14.0.

### Tissue culture

CMT-93 cells (CCL-223, ATCC), adherent HEK293 (CRL-1573, ATCC) cells were grown in high-glucose DMEM (4 mM L-glutamine, 4500 g/L glucose, 1 mM sodium pyruvate, and 1500 mg/L sodium bicarbonate) supplemented with 10% FBS. Expi293 cells (Thermo) were grown in Expi293™ Expression Medium (Thermo, A1435102). Cells were authenticated using STR profiling service from ATCC and tested for mycoplasma using Applied Biosystems MycoSEQ detection system based on qPCR. Media was changed every two days. HEK293 and Expi293 cells were transfected using X- tremeGENE9 (Roche) with a pCMV6 mammalian expression vector carrying a full-length *Dmbt1* gene (Origene MR212101). Cells were analysed 3 days post-transfection by staining with recombinant anti-HA SIgA labelled with alexa fluor 647 for imaging or with a rat anti-mouse DMBT1/GP340 antibody (clone 548031) for flow cytometry. The CMT-93 cells were used between passage number 5 and 20 post-ATCC handling. Treatments with antibodies were initiated immediately upon plating cells and were kept in media for five days, with one media change. At the endpoint, cells were counted using a flow cytometer analyser with counting beads and a live/dead stain (which we found to be more reproducible than manual or automated counting of trypan blue). CRISPR knockouts were performed using the lentiCRISPR system as previously described^71^. Briefly, guide RNAs were cloned into lentiCRISPR v2 plasmid (addgene #52961), adherent HEK293 cells were transfected using X- tremeGENE9 (Roche) reagent with individual lentiCRISPR plasmids and packaging vectors (VSV-G and Delta-VPR) to produce virus particle-containing supernatants. CMT-93 cells at 30-50% confluence were infected with individual viral supernatant by centrifuging for 90 minutes at 2200 rpm at room temperature and then incubating for 24 hours at 37C. Cells were selected with 5 µg/ml puromycin for the following 24 hours, after which they were provided with fresh media and treatment was initiated. RNA was isolated from tissue-cultured cells using Trizol and Qiagen RNeasy kits as described above (bulk RNA-Seq), first-strand cDNA synthesis was performed using the BioRad iScript kit, and quantitative real-time PCR was performed using Power SYBR Green PCR Master Mix (Applied Biosystems) on a QuantStudio 3 (Applied Biosystems). All primers and guide RNAs are listed in **Supplementary Data 5**.

### IgA co-purification and mass spectrometry

Colon organoids derived from IgA knockout mice, *ApcΔ/Δ*;*Kras^G12D/+^*;*Trp53Δ/Δ* (AKP)^72^ and metastatic organoids (both generated in Omer Yilmaz’s laboratory) were cultured for use in cellular lysate preparation. The IgA knockout mouse-derived organoids were isolated as previously described in this methods section, but this time cultured with IntestiCult™ Organoid Growth Medium (Mouse) (#06005 STEMCELL) supplemented with 1% Pen/Strep (Gibco 15140). After isolation, the organoids were maintained for two passages by resuspending them in cold media mixed with Matrigel (Fisher Scientific CB40230C) at a 1:1 ratio. In each passage, five 40 µL droplets of the cell suspension were plated in a 6- well plate, incubated at 37°C for 20 minutes, and then 3 mL of pre-warmed media was added. AKP and metastatic organoids were cultured in ADF+ media, composed of Advanced DMEM F12 (Invitrogen) supplemented with 5% heat-inactivated FBS (Sigma-Aldrich), 1% Penicillin-Streptomycin-Glutamine (Invitrogen), and 1% HEPES (Invitrogen). These organoids were plated in droplets, following the same procedure as for the mouse-derived organoid passages. To prepare the cells for lysis, all organoids were dissociated in cold ADF+ media to break down the Matrigel. The cell suspensions were centrifuged at 1000 rpm for 5 minutes. After removing the supernatant, 1 mL of TrypLE (Thermo Fisher Scientific 12605010) was added to the cell pellet, followed by incubation in a 37°C water bath for 25 minutes. To stop the dissociation and wash the cells, 5 mL of ADF+ media was added to each organoid type. Cells were then pelleted again by centrifugation at 1500 rpm, and the supernatant was carefully removed. For the pulldown, cells were lysed using a lysis buffer consisting of 0.50% Igepal CA-630 in 1x PBS supplemented with a protease inhibitor cocktail (Roche). The cell pellet was resuspended in 500 µL of this lysis buffer and briefly vortexed. The lysate was incubated on ice for 30 minutes to ensure complete lysis and then centrifuged at 14,000 g for 10 minutes at 4°C. Meanwhile, Pierce^TM^ Protein L Magnetic Beads (Thermo Scientific) were washed by adding 100 µL of beads to 300 µL of PBS-T in six tubes.

The beads were gently vortexed for 1 minute and collected using a magnet. After removing the supernatant, the beads were washed again with 1 mL of PBS-T, followed by gentle vortexing and removal of the supernatant. The resulting lysis supernatant of each organoid was transferred to two different tubes containing the washed beads - 240 µL of supernatant added to each tube. The division was made to achieve two conditions: one that received the faecal SIgA and one negative control, where the supernatant was incubated with beads in the absence of SIgA. Next, 100 µL of SIgA (at a concentration of 600 µg/mL) was introduced into each designated tube, and the positive and negative mixtures were incubated for 2 hours at 4°C with gentle agitation. The beads were then collected using a magnet, and the supernatant was removed. The beads were washed three times with 200 µL of PBS-T, vortexed briefly and carefully between washes, and collected using a magnet to remove the supernatant each time. Finally, 22.5 µL of elution buffer (0.1M glycine at pH 2-2.5) was added to the beads, and the mixture was incubated at room temperature for 10 minutes with occasional vortexing. The eluate was combined with 2.5 µL of 1M Tris-HCl at pH 8. Proteins bound to magnetic beads were eluted through partial trypsin digestion (Promega). The supernatants were reduced with DTT, alkylated using iodoacetamide, and subjected to overnight redigestion with trypsin and Endopeptidase LysC (Wako/Fuji). Digestion was halted with TFA, followed by peptide cleanup via solid-phase extraction (SPE’ed). The peptides were then analyzed by LC-MS/MS, utilizing a 70-minute analytical gradient (2%B to 38%B) on a 12 cm built-in emitter column with high resolution and mass accuracy (EasyLC1200 and Lumos Fusion, Thermo Fisher). The resulting data was searched against the UniProt mouse database, concatenated with common contaminants, using Proteome Discoverer (v1.4.1.14) and MASCOT (v2.8.01). Data were filtered with a Percolator-calculated false discovery rate (FDR) of 1% or better. Further processing was done by filtering peptide-spectrum match (PSM) counts for each protein and organizing them into a count matrix. Missing values were imputed as zero. The matrix was used to generate a DESeq2 object, and results were obtained without fold change (FC) shrinkage. Differential analysis was performed to identify significant changes in protein abundance between IgA+ and IgA- groups. Results were visualized using a volcano plot, where proteins with Log2FC ≥ 1 or ≤ −1 and adjusted p-values < 0.05 were considered for downstream analysis.

### Faecal Immunoglobulin quantification by ELISA

Faeces were weighed and dissolved in 1 mL PBS with 0.1% tween-20 (Sigma-Aldrich, P9416) and centrifuged at 1,000 g for 5 minutes at 4°C. Supernatant was transferred into a new tube and centrifuged at 16,000 g for 10 minutes at 4°C. This supernatant was stored at −20°C until ELISA measurements. ELISA high binding half area plates (Greiner Bio-One, 675061) were coated with antibodies against the respective isotype (SouthernBiotech, IgA: 1040-01, IgG: 1030-01, IgM: 1020-01) 1:1000 in PBS at 4°C overnight. The following day, plates were blocked with 2% BSA in PBS for 2 hours at room temperature, washed with PBS containing 0.1% tween-20, and incubated with the faecal Ig extracts and standard dilutions for each isotype for 2 hours at room temperature. Plates were washed with PBS containing 0.1% tween-20 and incubated with antibodies against the respective isotype coupled to HRP (SouthernBiotech, IgA: 1040-05, IgG: 1030-05, IgM: 1020-05) for 1 hour at room temperature. After an additional two washed with PBS containing 0.1 % tween-20 and two washes with PBS, plates were incubated with 3,3’,5,5’-Tetramethylbenzidine (Sigma-Aldrich, T0565) at room temperature until the standard curves showed a good resolution of colour gradients upon which the reaction was stopped with 1M Hydrchloric acid (VWR, 450550025). Absorption was read at 450 nm and 600 nm on a Agilent Biotek Synergy HTX (using Gen5 software).

### Isolation of plasma cells from mouse colon (lamina propria)

Mice were sacrificed by cervical dislocation and the colon was harvested as previously described (see ‘Colon Epithelial Cell Isolation’). The tissue was cut into 1 cm pieces and incubated in PBS with 1 mM DTT (Thermofisher Scientific) for 10 minutes at RT on a rolling shaker, followed by a brief shake to remove mucus. The tissue was transferred twice to RPMI with 30 mM EDTA and incubated for 10 minutes at 37 C on rolling shaker. For collection of lamina propria lymphocytes, the supernatant was discarded and the tissue transferred to gentleMACS C tubes (Miltenyi Biotech) with digestion mix containing Collagenase VIII (0.05 mg/ml, Sigma Millipore), 0.25 mg/ml trypsin inhibitor (Roche), and DNAse I (0.15mg/ml, Roche) in RPMI 1640 (Gibco) supplemented with 2% FCS, 10 mM HEPES, and 1 mM sodium pyruvate. After tissue digestion by gentleMACS Octo Dissociator (Miltenyi Biotech) according to manufacturer’s instructions, cells were filtered through a 70 µm strainer, washed and stained in FACS buffer for quantification of plasma cells by flow cytometry. For surface staining the following antibodies were used: B220 (RA3-6B2, BD Bioscience), CD3 (17A2, BioLegend), CD45 (281-2, BioLegend), CD98 (RL388, BioLegend), CD138 (281-2, BioLegend), DMBT1 (548031, R&D Systems), F4/80 (BM8, eBioscience), IgA (mA-6E1, Invitrogen), NK1.1 (PK136, Invitrogen) and Zombie NIR Fixable Viability Kit (BioLegend). Data was collected on a FACS Symphony A5 (BD Bioscience) and analyzed by FlowJo 10.8.2 software.

### Microscale electrophoresis of purified DMBT1 protein and antibodies

Human sIgA from breast milk (Athens, 16-13-090701) and recombinantly produced human dIgA, IgA and IgG1 were labelled using the Protein labelling Kit RED-NHS 2nd Generation (NanoTemper, MO- L011) according to the manufacturer’s instructions except for removing excess dye which was done using Zeba spin columns (Thermo Scientific, 89877) according to their respective instructions. After verifying adequate labelling, we performed binding assays of each labelled antibody against human Dmbt1 (R&D systems, 11032-DB) at dilutions ranging from 2.78 µM to 8.47E-05 µM using Premium Capillaries (NanoTemper, MO-K025) on a Monolith NT.115 Microscale Thermophoresis machine (NanoTemper). Data was processed using the NanoTemper Monolith software v.X.

### Statistical methods

Unless otherwise stated, mean and standard error are plotted for all figures. For comparisons between two groups, two-tailed unpaired t-tests were performed. For three or more groups, ordinary one-way ANOVA tests were performed with Dunnett’s correction for multiple comparisons to a single control group or Tukey’s correction for multiple comparisons amongst all groups. For experiments with repeated measures over time, repeated-measure 2-way ANOVA tests were performed. Exact p values are reported on graphs when the p value or adjusted p value is less than 0.1. Exact p values are not calculated below 0.0001 and are simply reported as such. For sequencing analyses, application-specific statistical methods are described in the respective methods sections or figure legends.

## Data Availability

All raw sequencing data for bulk RNA-Seq, 16S rRNA amplicon sequencing, and single cell RNA-Seq are available on the NCBI Sequence Read Archive (SRA) under a unified project: PRJNA1023250. The following freely-accessible databases were also used: UniProt (uniprot.org) Tabula muris and Tabula sapiens (biohub.org), and GSEA gene sets (gsea-msigdb.org).

## Acknowledgements

The authors thank J. Faith (Mt. Sinai), D. Wesemann (Brigham and Women’s), M. Koch (Fred Hutchinson), N. Palm (Yale), L. Mesin (Rockefeller), and members of the Mucida and Victora labs at Rockefeller University for useful discussions. S. Leonard assisted with protein purifications at University of Illinois. Germ-free mice were generated and maintained by A. Rogoz at Rockefeller University. *Dmbt1* transmembrane-knockout mice were generated by J. Bortolatto at Rockefeller University. All animals were monitored and cared for by the staff of the Comparative Bioscience Center at Rockefeller University. Sequencing was performed at the Genomics Resource Center of Rockefeller University (RRID:SCR_020986). Proteomics data were generated by the Proteomics Resource Center at Rockefeller University (RRID:SCR_017797) with assistance from H. Molina using instrumentation funded by the Sohn Conferences Foundation and the Leona M. and Harry B. Helmsley Charitable Trust. Microscale electrophoresis was performed at the Fisher Drug Discovery Resource Center at Rockefeller University (RRID:SCR_020985) with assistance from J. Fraser Glickman.

## Author contributions

G.P.D. and D.M. conceived and initiated the project. G.P.D. designed and performed most of the experiments, with guidance from D.M.. G.L.R. performed organoid experiments, purified IgA, performed the pull-down assay, and assisted with most of the experiments. M.S. performed the colonoscopy procedures and cultured organoids. C.W. performed tissue culture and biochemical binding experiments. G.P.D. analyzed 16S rRNA amplicon sequencing, I.M. analyzed single-cell RNA-Seq, and G.C. analyzed bulk RNA-Seq, with guidance and analysis oversight from T.B.R.C. N.L.D. assisted with flow cytometry and tissue culture experiments. C.H. performed plasma cell flow cytometry and assisted with fluorescence imaging. D.Z. and B.M.S. designed and produced recombinant secretory component and IgA variants. G.P.D., B.A., and B.S.R designed and executed epithelial cell sorting and single-cell transcriptomics. S.G. and B.S.R. advised and assisted with mouse models of colorectal cancer. E.G.L. and S.C.E. provided the 2025 IgA knockout mouse and designed and performed cancer experiments. K.B., S.F.T., B.M.S., G.D.V., and D.M. supervised, guided experimental design, and directed interpretation of results. G.P.D. and D.M. wrote the manuscript, and all authors edited the manuscript.

---

**Extended Data Fig. 1.**
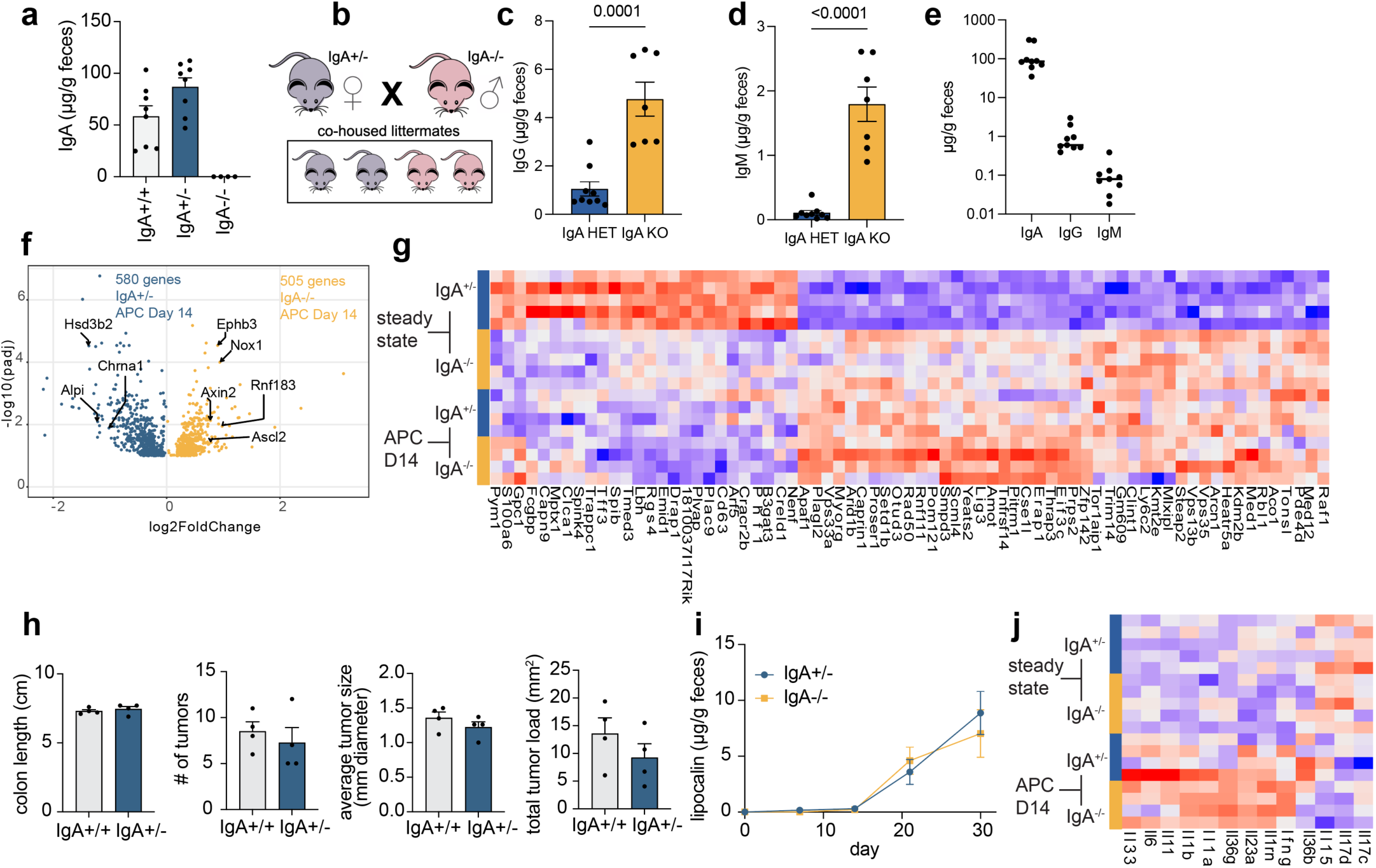
Additional data related to IgA knockout model and APC-driven colorectal cancer model. **a**, ELISA of faecal IgA in homozygotes and heterozygotes of the IgA^−/−^ strain crossed with C57BL6/J (n = 8, 8, 4). **b**, Breeding scheme for littermate controls. **c**, ELISA of faecal IgG and **d**, IgM in littermates from this breeding scheme (n = 9,7). **e**, Combined visualization of faecal antibody isotype ELISA data from the IgA^+/−^ mice in c-d (n = 9, statistics not used). **f**, Differentially expressed genes in bulk RNA-Seq from mid-colon tissue of IgA^+/−^ and IgA^−/−^ littermates fourteen days after inducing colon epithelial-specific loss of *APC* (n = 4). **g**, Set of genes identified as modulated by both *APC* loss (steady state vs. APC Day 14 comparison) and IgA deficiency at steady state (IgA^+/−^ vs. IgA^−/−^). Interestingly, the direction of the effect of IgA deficiency is toward the direction caused by *APC* loss (n = 5, 5, 4, 4). **h**, Endpoint analyses of the APC-driven cancer model using IgA^+/+^ and IgA^+/−^ littermates (n = 4). **i**, ELISA for faecal lipocalin in the APC-driven cancer model (n = 3, from one cohort, representative, repeated measure 2-way ANOVA not significant). **j**, Cytokine gene expression in the bulk colon tissue RNA-Seq shows no IgA-dependent alterations in inflammatory pathways (n = 5, 5, 4, 4) (mean and SEM, t-tests unless otherwise stated, p values hidden when p > 0.1).

**Extended Data Fig. 2.**
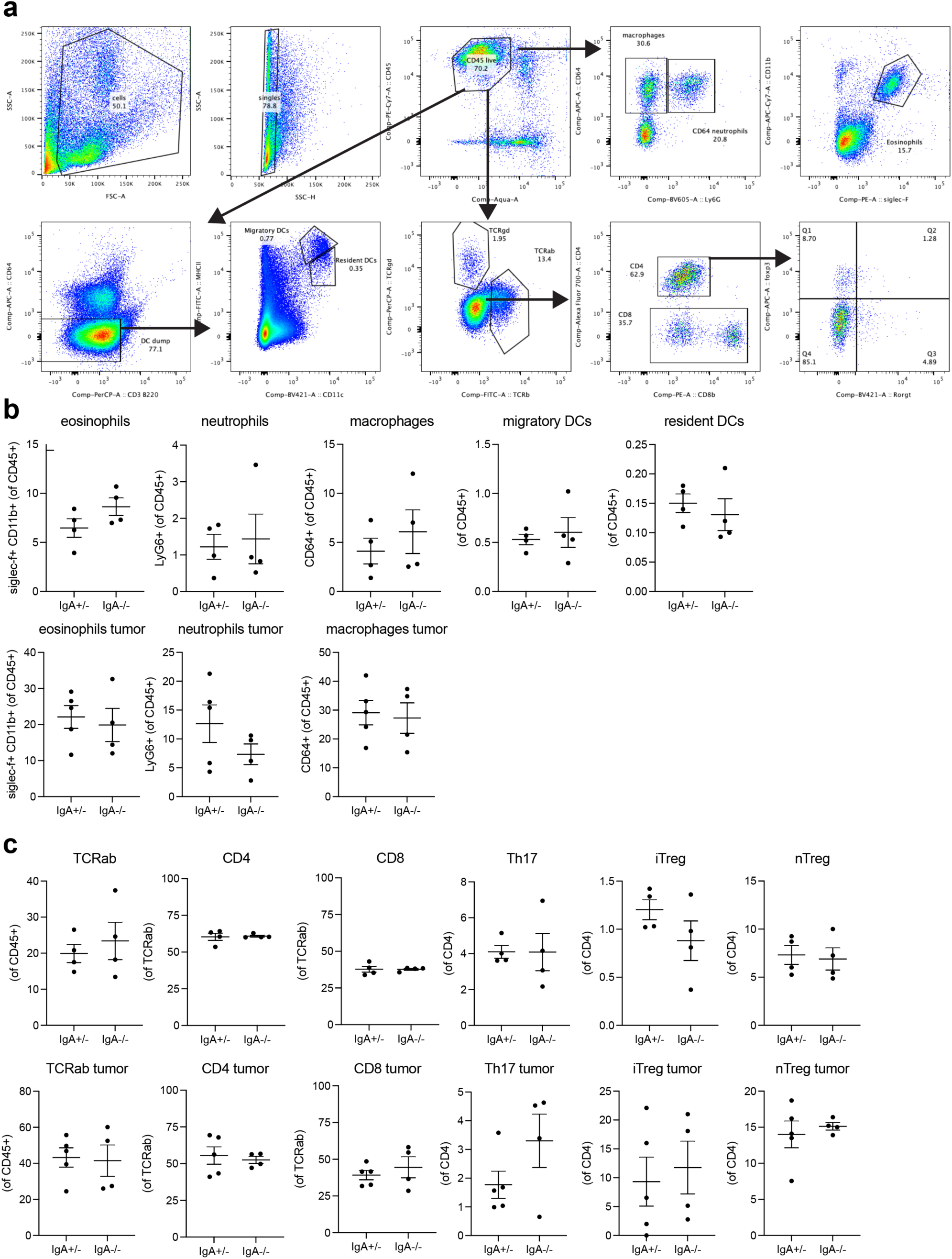
Immune profiling of normal colon lamina propria and tumour-containing tissue shows no IgA-dependent differences in co-housed littermates. **a**, Gating strategy for innate and adaptive immune profiling of major colon-relevant subpopulations. **b**, Innate and **c**, adaptive cell populations in colon lamina propria (top row) or APC mutation-induced endogenous tumours (bottom row) isolated from IgA^+/−^ and IgA^−/−^ littermates (n = 4, pooled two independent experiments for healthy mice and n = 5, 4, pooled two independent experiments for tumours) (mean and SEM, t-tests, p values hidden when p > 0.1).

**Extended Data Fig. 3.**
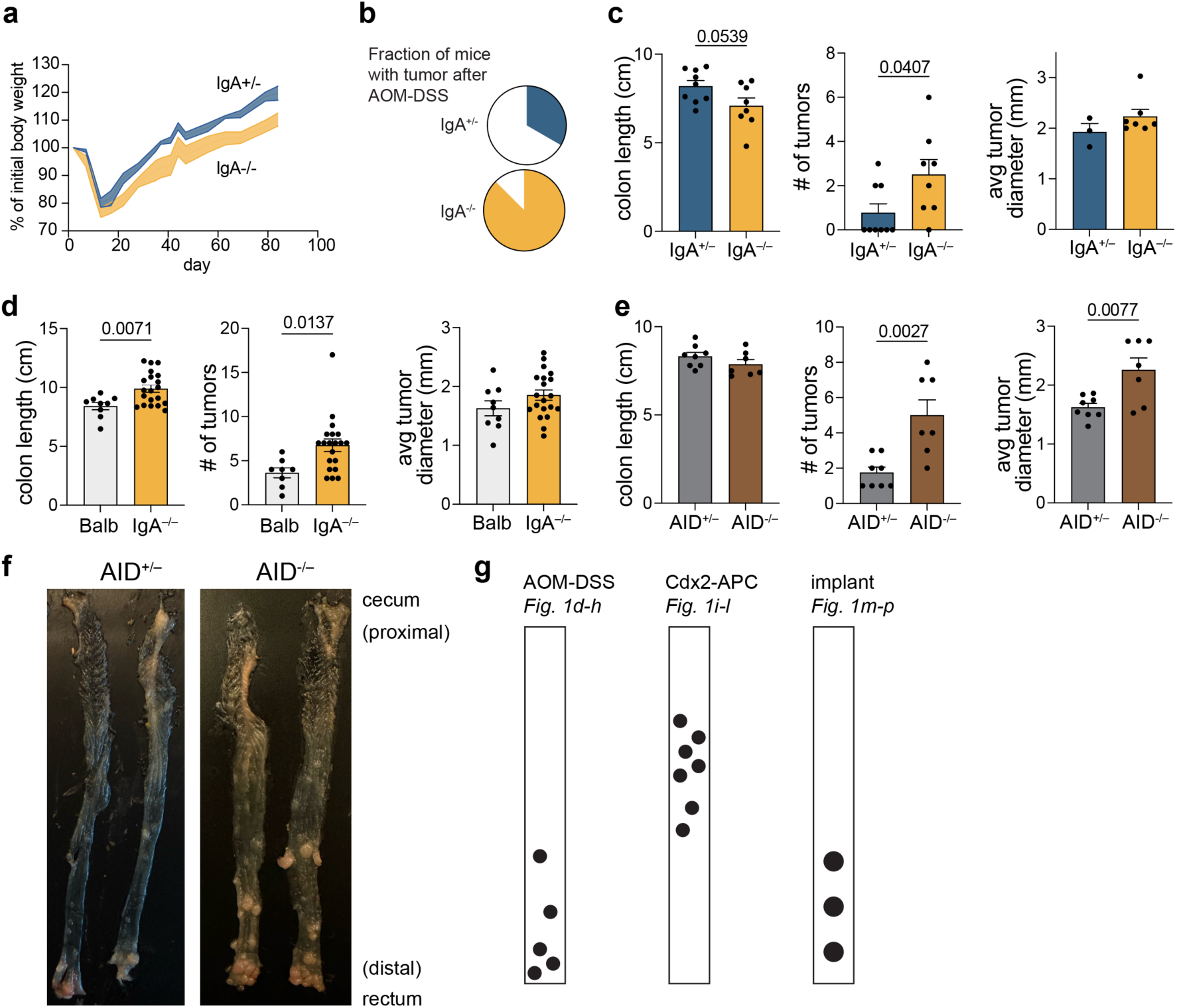
Additional data related to different IgA deficiency models and colon cancer models. **a**, Body weight change during AOM-DSS (n = 4, from one cohort, representative). **b**, Tumour frequency in the AOM-DSS cancer model (n = 9, 8) (t-tests unless otherwise stated, p values hidden when p > 0.1). **c-e**, Colon length, tumour count, and average tumour size outcomes in AOM-DSS for 1999 IgA^−/−^ mice (**c**, as in Fig. 1j, pooled 2 cohorts, n = 9, 8 except mice without tumours excluded from average tumour size figure: n = 3, 7), 2025 BALB/c IgA^−/−^ mice (**d**, as in Fig. 1k, pooled 2 cohorts, n = 9, 20), and AID^−/−^ mice (**e**, as in Fig. 1l, representative of 2 cohorts, n = 8, 7). **f**, representative images of colons with tumours from AID^−/−^ mice at endpoint of AOM-DSS (representative of 2 cohorts, n = 8, 7). **g**, Schematic showing differences in general tumour localization in the three colorectal cancer models used in this work. (mean and SEM, t-tests, p values hidden when p > 0.1)

**Extended Data Fig. 4.**
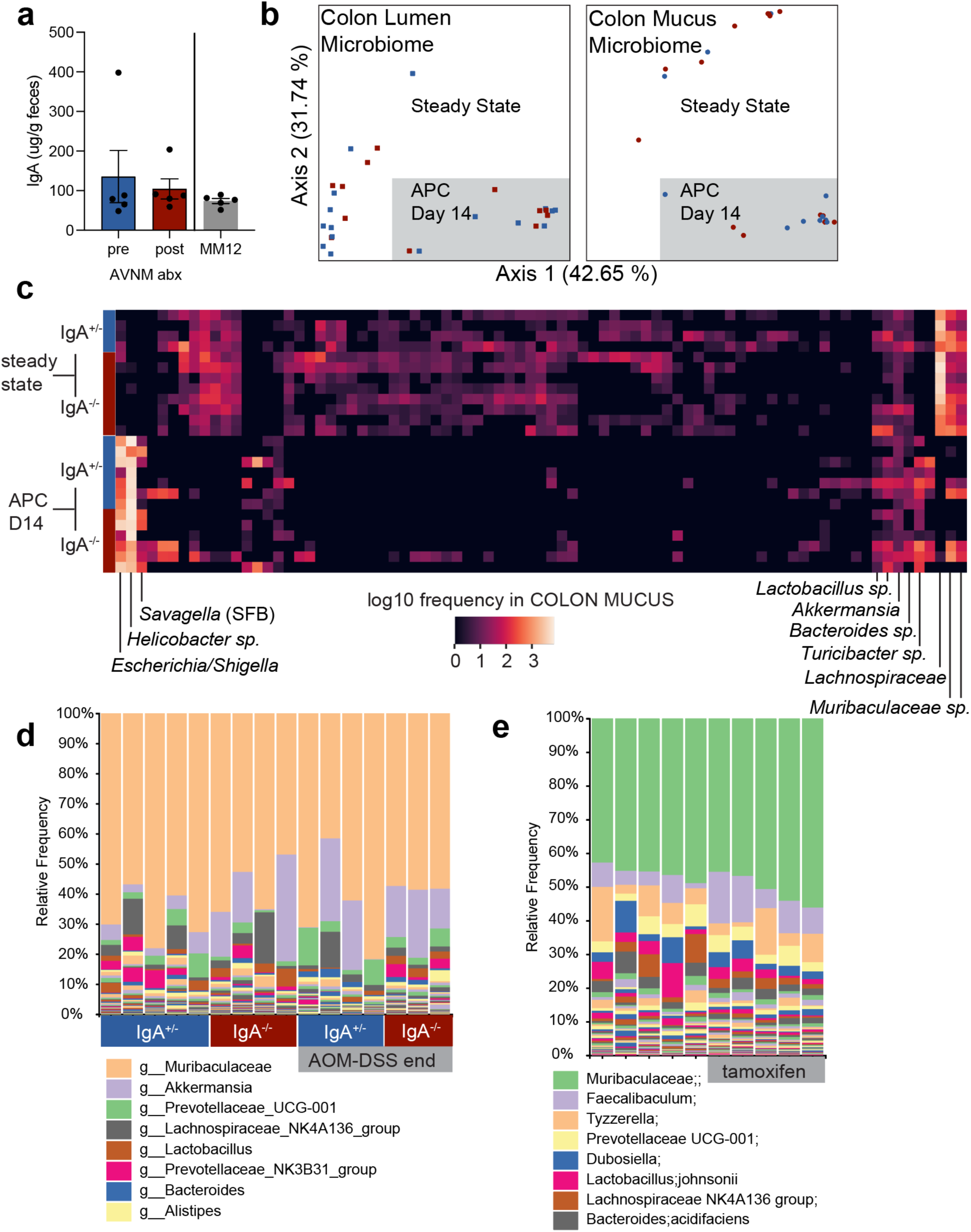
Microbiome profiling data. **a**, ELISA for IgA in faeces of IgA^+/−^ mice before and after antibiotics treatment as in Fig. 2b and in mice colonized with oligo-mm12 as in Fig. 2e (n = 5) (mean and SEM, one-way ANOVA, p values hidden when p > 0.1). **b**, Beta diversity (weighted unifrac distance) of microbiome measured by 16S rRNA amplicon sequencing of samples from IgA^+/−^ (blue) and IgA^−/−^ (red) mice showing no genotype-specific alterations (lumen n = 9, 6, 7, 6, mucus n = 4, 8, 7, 6). **c,** Heatmap of amplicon sequence variants (columns) identified in colon mucus samples from indicated mice (rows) demonstrates dramatic shift caused by *APC* loss, but no effect of IgA. **d**, Relative abundance of ASVs identified by 16S rRNA amplicon sequencing of faeces from IgA^+/−^ and IgA^−/−^ littermates before and after AOM-DSS treatment (n = 5, 4, 4, 3). **e,** Relative abundance of ASVs identified by 16S rRNA amplicon sequencing of faeces from wildtype C57BL/6j mice before and two weeks after a two-dose tamoxifen treatment that matches the induction protocol used to knockout APC in Fig. 1d-h demonstrates that the microbiome shift observed is not due to tamoxifen injection (n = 5).

**Extended Data Fig. 5.**
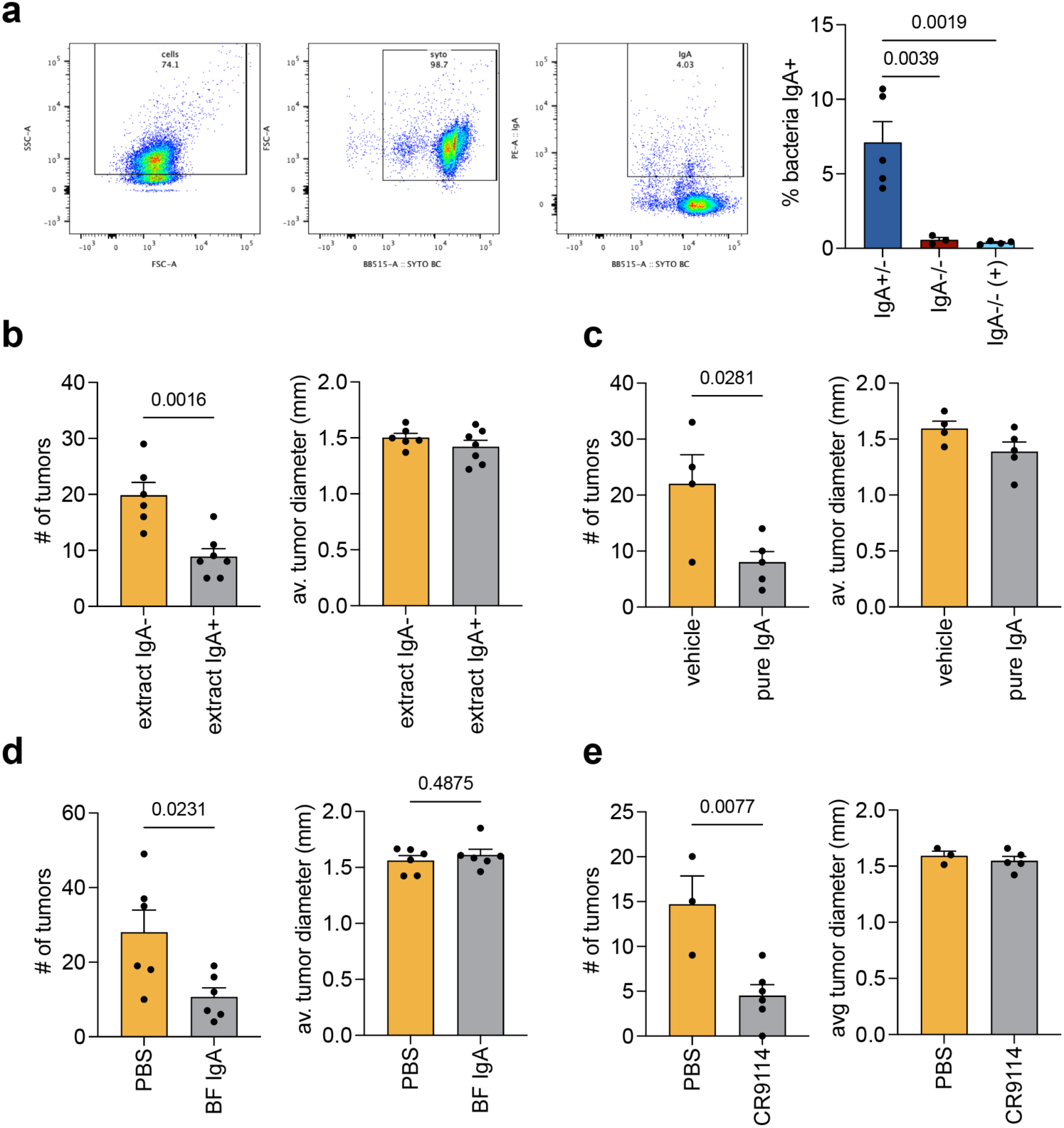
Additional results related to IgA gavage treatments. **a**, Gating strategy for flow cytometry for IgA coating of bacterial cells in faecal samples and summary data showing a lack of IgA binding to bacteria when IgA^−/−^ mice were gavaged with faecal IgA purified from germfree mice monocolonized with *Bacteroides fragilis* (n = 5, 3, 4). **b-e**, Counts and sizes of tumours from IgA^−/−^ mice treated with **b**, faecal extracts (n = 6, 7), **c**, purified faecal IgA from SPF mice (n = 4, 5), **d**, purified faecal IgA from mice mono-colonized with *Bacteroides fragilis* (n = 6), or **e**, recombinant CR9114 anti-influenza HA monoclonal secretory IgA (n = 3, 5) (mean and SEM, t-tests or one-way ANOVA, p values hidden when p > 0.1).

**Extended Data Fig. 6.**
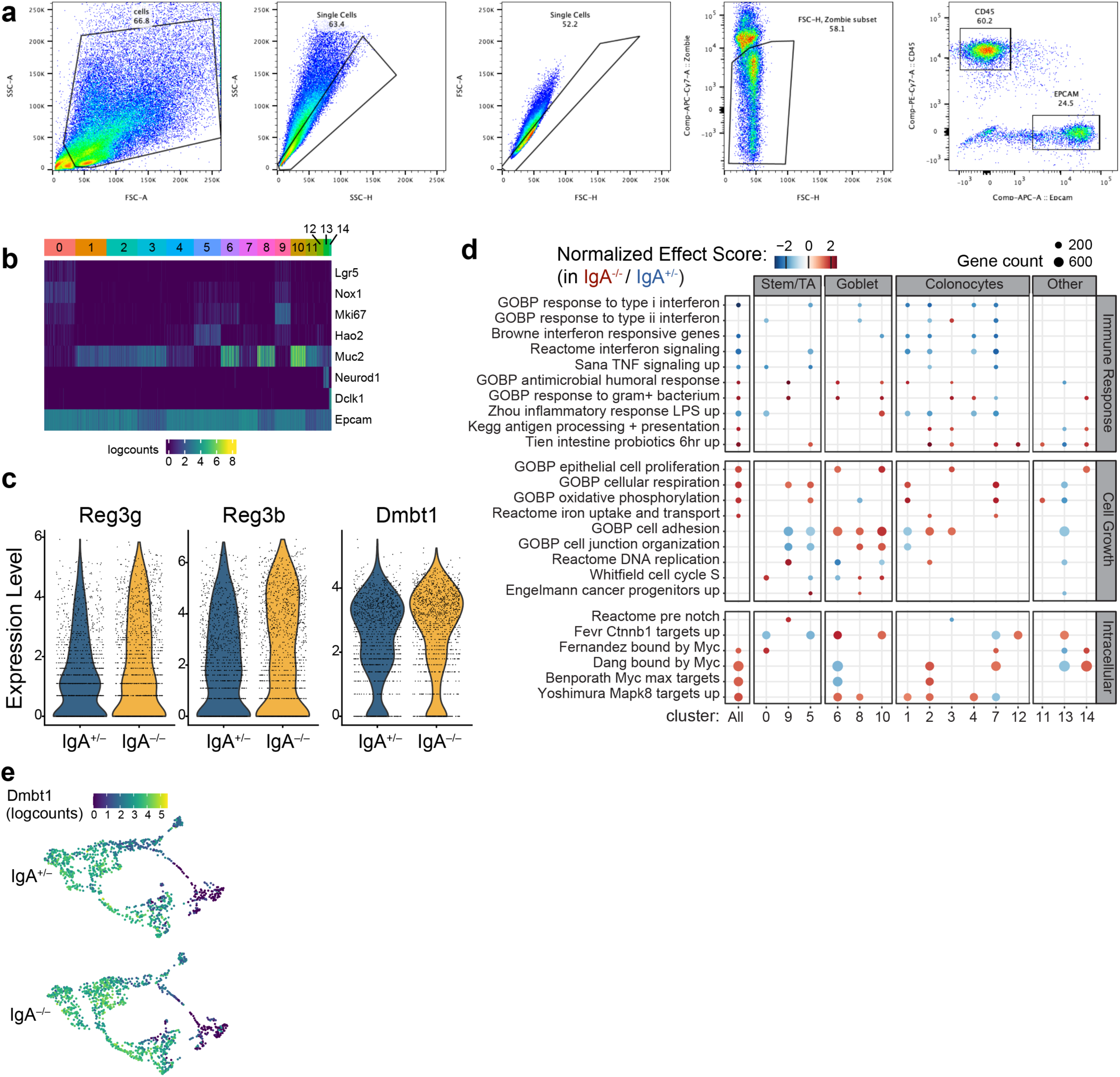
Additional data related to colon epithelial single cell RNA-Seq. **a**, Gating strategy for sorting live EPCAM+, CD45- epithelial cells for single-cell transcriptomics (n = 3). **b**, Various markers to aid in identifying clusters: *Lgr5* (stem), *Nox1* (transit-amplifying), *Mki67* (cycling), *Hao2* (immature), *Muc2* (low: mature, high: goblet), *Neurod1* (enteroendocrine), *Dclk1* (Tuft), and *Epcam* (pan-epithelial). **c**, Expression level of *Reg3g, Reg3b,* and *Dmbt1* in all cells. **d**, Full version of Fig. 3g with all clusters: Gene set enrichment analyses in individual clusters based on IgA genotype. Dots are visible where adjusted p value < 0.05, size of dot represents the number of genes int he pathway, and color represents the normalized enrichment score of that pathway (IgA^−/−^-enriched is positive, red). **e,** *Dmbt1* reads across cell types.

**Extended Data Fig. 7.**
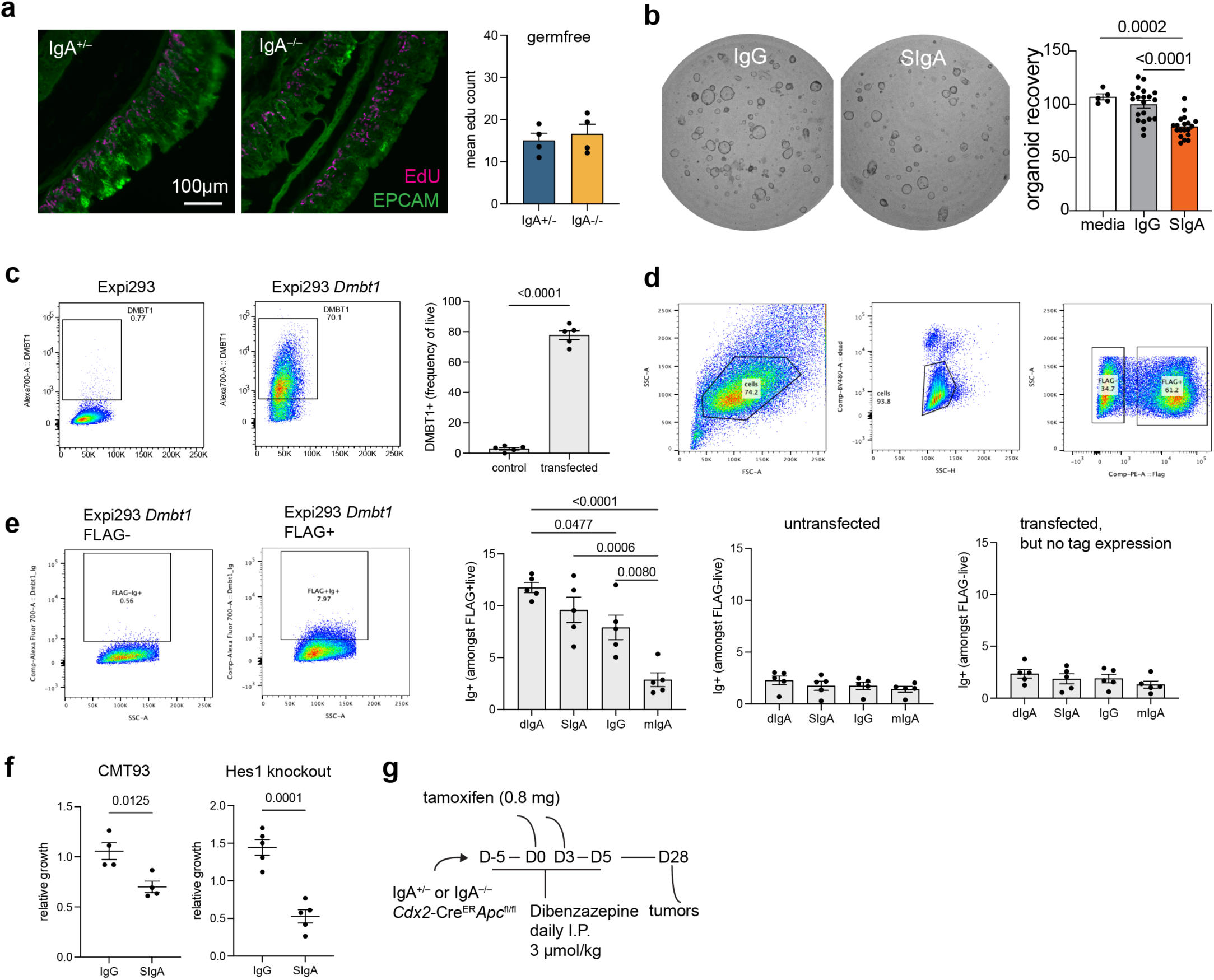
Additional data related to epithelial proliferation. **a**, EdU (pink) and EPCAM (green) staining in germfree mice after two injections (−24 hours and −4 hours) of EdU showed no difference between IgA^+/−^ and IgA^−/−^ littermates (n = 4). **b**, Primary mouse colon organoids were cultured from isolated crypts in the presence of indicated antibody isotypes imaged by brightfield microscopy. Organoid counting from four independent experiments pooled and normalized to the IgG treatment control (n = 5, 20, 19). **c**, Flow cytometry of surface DMBT1 on Expi293 cells: wildtype (left) and expressing a membrane isoform of *Dmbt1* (right), with summary data graphed (n = 5 independent transfections). **d**, Gating strategy for transfected Expi293 cells that are live and have a FLAG tag marking transgene expression. **e**, Binding of Expi293 cells to different antibody structures was dependent on transfection with the *Dmbt1* expression construct and also dependent on FLAG tag expression marking surface DMBT1 protein (n = 5 independent transfections performed over 3 independent experiments). **f**, SIgA regulation of CMT93 cell growth was not dependent on *Hes1* gene (n = 4, 4, 5, 5). **g**, Experimental design for Notch inhibition to suppress epithelial proliferation in the context of IgA deficiency and *APC* loss (mean and SEM, t-tests or one-way ANOVA, p values hidden when p > 0.1).

**Extended Data Fig. 8.**
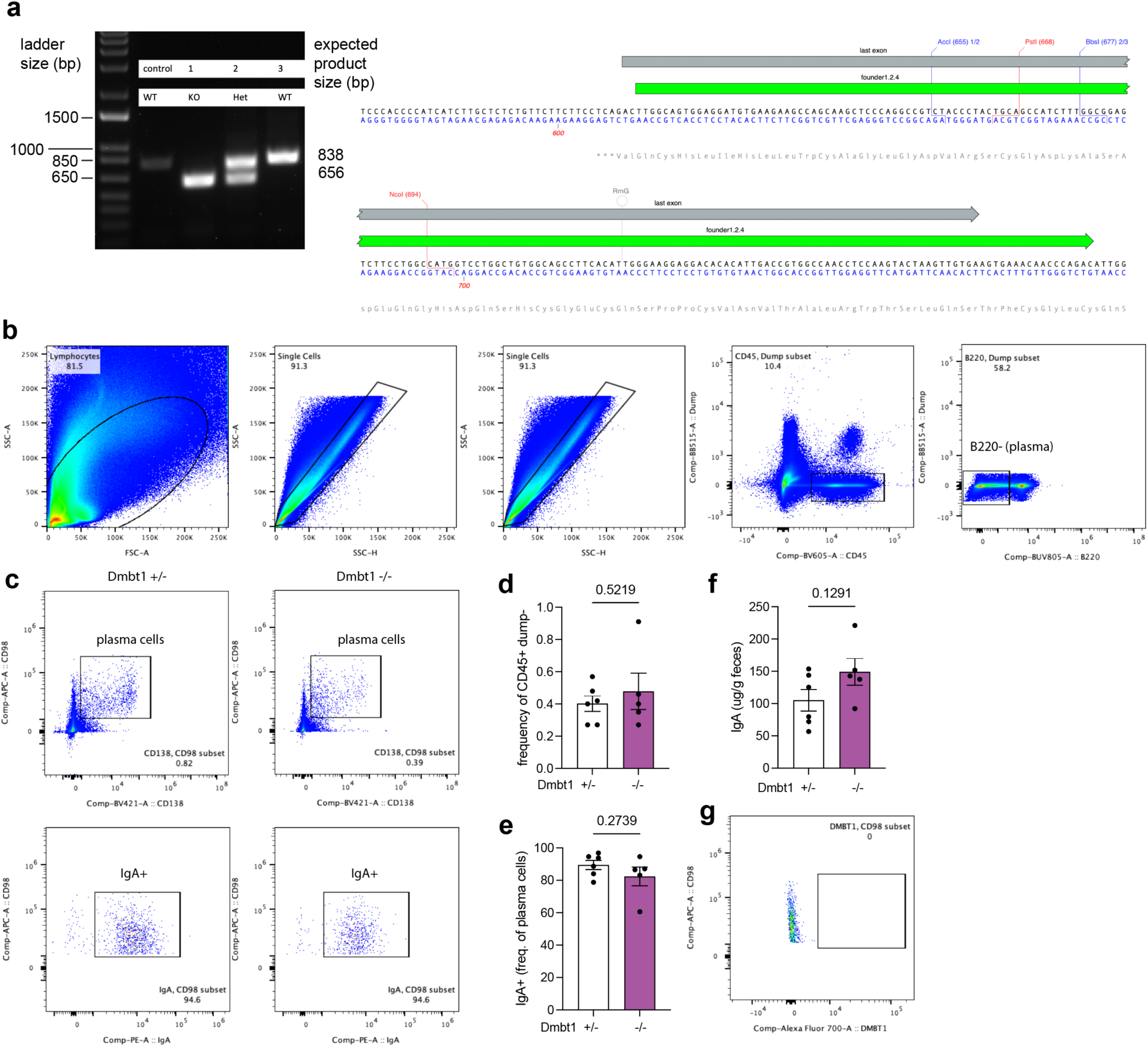
Additional data regarding transmembrane *Dmbt1* knockout mouse model. **a**, DNA gel showing knockout of 182 basepair in the last exon of mouse *Dmbt1* and the sequence-confirmed knockout on mouse genome (green track) (original gel of two individual founder mice). **b**, Gating strategy for plasma cells in the colon lamina propria. **c,** Total plasma cells and IgA+ (bottom row) plasma cells in *Dmbt1^+/−^* and *Dmbt1^−/−^* littermate mice. **d**, Quantification of total plasma cells and **e**, IgA+ plasma cells (n = 6, 5, pooled two independent experiments). **f,** Faecal IgA levels quantified by ELISA in *Dmbt1^+/−^* and *Dmbt1^−/−^* littermate mice (2 cohorts pooled, n = 6, 5). **g**, Colon lamina propria plasma cells do not express DMBT1 (mean and SEM, t-tests).

**Extended Data Fig. 9.**
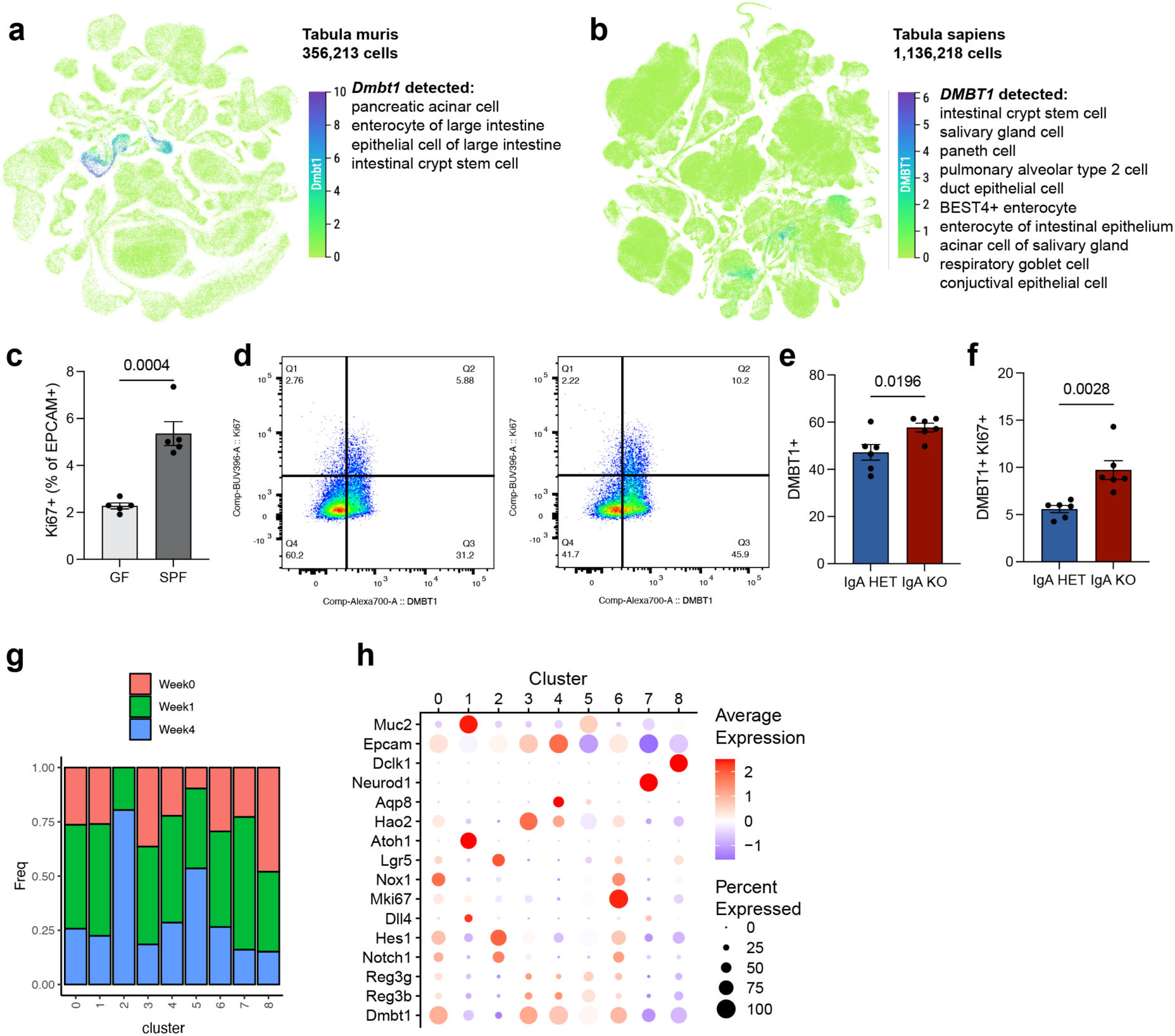
Additional data related to DMBT1 association with physiological epithelial proliferation and decoupling of DMBT1 and Notch during tumourigenesis. Single-cell expression of *Dmbt1* across mouse tissues (**a**) and human tissues (**b**) reveals remarkable conservation and specificity to epithelial cell subsets. **c**, Plot of proliferating epithelial cells to directly compare germfree and SPF mice from Fig. 5a (n = 5). **d**, Flow cytometry analysis of DMBT1 and Ki67 in colon epithelial cells isolated from IgA^+/−^ and IgA^−/−^ littermate animals demonstrates **e**, increased DMBT1 in IgA^−/−^ mice, consistent with single cell RNA-Seq in Fig. 3a-f (pooled two cohort, n = 6). **f**, Increased proliferation of epithelial cells in IgA^−/−^ mice occurs amongst cells that are DMBT1+ (same mice as e). **g**, Relative contribution of each timepoint to clusters in single cell RNA-Seq analysis in Fig. 5. Cluster 2, absent prior to APC knockout, represents the transformed tumour cells. **h**, Differential expression across clusters (Fig. 5e), reveals that the tumour cells (cluster 2) lose epithelial identity and dramatically lose *Dmbt1* expression. (mean and SEM, t-tests, p values hidden when p > 0.1)

**Extended Data Table 1.** Microscale electrophoresis of purified DMBT1 protein and antibodies. Purified human DMBT1 protein and human antibodies were used for microscale electrophoresis binding assays, revealing low affinity interactions between DMBT1 and different antibody structures. Statistics reported within table.

|  | dimeric IgA | monomeric IgA | IgG | Secretory IgA |
| --- | --- | --- | --- | --- |
| N | 3 | 4 | 3 | 3 |
| Exc. power | 60% | 60% | 60% | 20% |
| MST power | 40% | 40% | 40% | 40% |
| Kd | 429 nM | 7.01 μM | 199 nM | 1.09 μM |
| Kd confidence | +/- 208 nM | +/- 7.95 μM | +/- 45 nM | +/- 1.50 μM |
| Amplitude | 5.197 | 19.349 | 17.379 | 1.626 |
| Signal to noise | 9.850 | 41.537 | 17.985 | 5.121 |

## Supplementary Information Guide

**Supplementary Figure 1:**
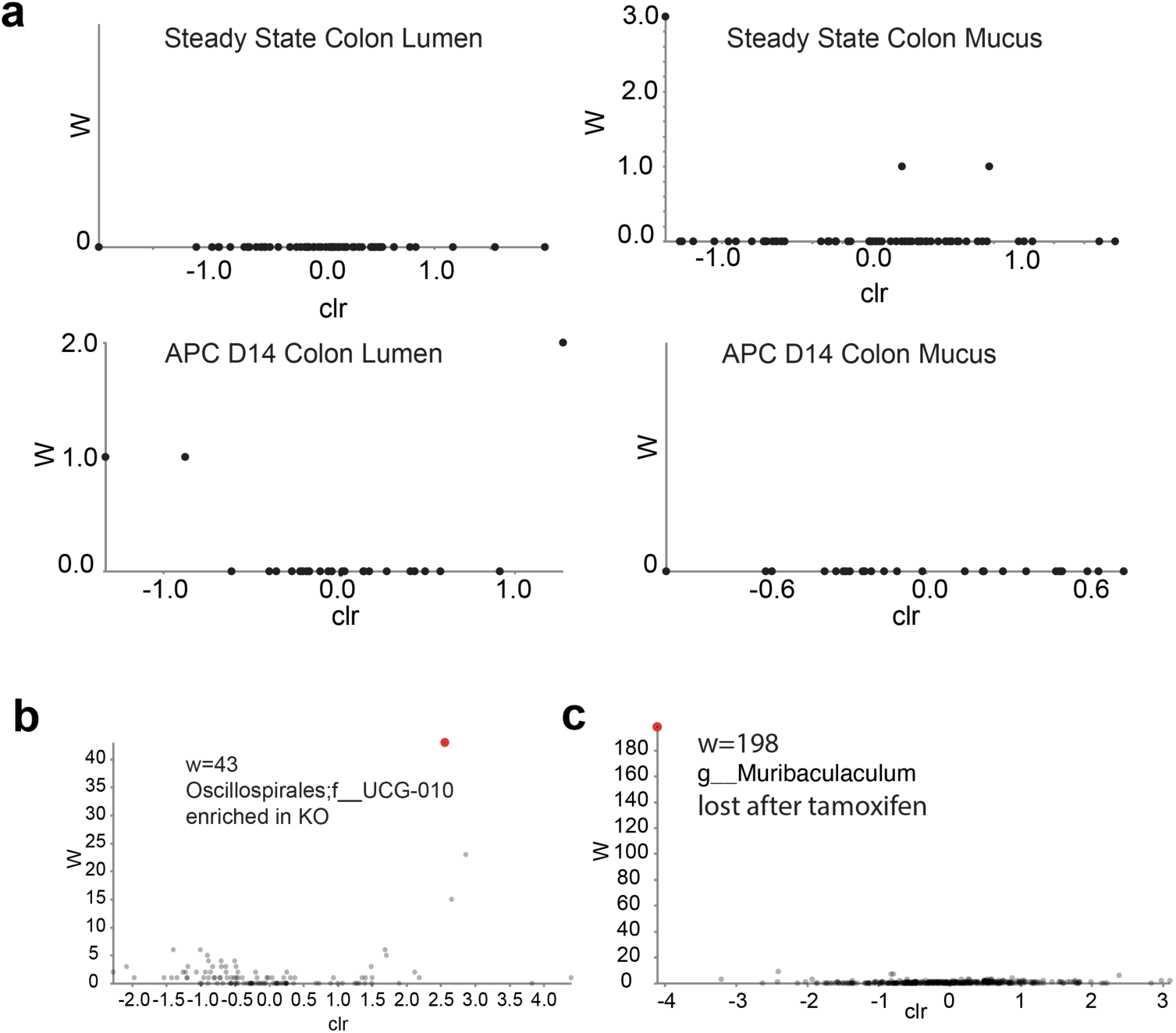
ANCOM statistical tests for microbiome data. **a**, ANCOM differential abundance testing of IgA^+/−^ vs. IgA^−/−^ littermates in the indicated sample types identified no IgA-dependent alterations in the microbiome (lumen n = 9, 6, 7, 6, mucus n = 4, 8, 7, 6, none significant by W statistic of ANCOM). **b**, Scatterplot shows results of ANCOM statistical test for differential abundance, identifying a single low-abundance ASV that is enriched in knockout mice following AOM-DSS (n = 5, 4, 4, 3). **c**, Scatterplot shows results of ANCOM statistical test for differential abundance, identifying a single low-abundance ASV that is lost upon tamoxifen treatment (n = 5).

**Supplementary Data: Steady state bulk RNASeq.** List of genes and associated statistics for those up and downregulated depending on IgA genotype during steady state (no cancer model), as shown in Figure 1.

**Supplementary Data 2: APC Day 14 bulk RNASeq.** List of genes and associated statistics for those up and downregulated depending on IgA genotype 2 weeks after loss of *APC* in epithelial cells (cancer model, pre- tumour), as shown in Figure 1.

**Supplementary Data 3: Steady state single cell RNASeq.** List of genes and associated statistics for those up and downregulated by IgA knockout (positive fold change = higher in IgA knockout) across all cells and within each cluster of the colon epithelial single cell RNA-Seq experiment in Figure 3.

**Supplementary Data 4: Mass Spec data of IgA pulldown from organoid lysates.** List of peptides and associated statistics for those enriched in pulldown with IgA (vs. control without IgA), showing mostly Ig- related peptides as expected, with a few notable hits (including identification of DMBT1).

**Supplementary Data 5: Primers and guide RNAs.** List of primers used in the study for PCR and guide RNAs for cell line and mouse knockouts.

## Supplementary Discussion

Although DMBT1 has been the subject of substantial research, its physiological functions and impact on cancer have been unclear. While the encoding gene was initially described as a possible tumour suppressor^73^, subsequent studies have revealed a much more complex relationship with cancer, reflecting tissue-specific biological functions. For instance, coating culture plates with hensin, a secreted form of DMBT1, is sufficient to induce ES cells to develop a columnar morphology^74^, reflecting potential biophysical functions of DMBT1 in cell shape and division. Consistent with this, DMBT1 is induced in Barrett’s metaplasia^75^ and is a general marker of gastric metaplasia^76,77^, both of which involve conversion to a columnar epithelial phenotype. DMBT1 appears to also have beneficial functions; in the case of head and neck squamous cell carcinoma, it may prevent invasiveness and metastasis^78^. DMBT1 may be involved in inflammation-driven metaplasia, reflecting aberrant antibacterial responses by epithelia, whereas at some point, it can be lost in adenomas, corresponding to the loss of epithelial identity and cell shape. In the context of colorectal cancer, *DMBT1* does not emerge in genetic studies as being frequently mutated. However, in an exhaustive proteomics analysis of colon polyp biopsies with a ten-year follow-up of patients, DMBT1 loss at the protein level was the overall best predictor of future progression to cancer^79^. This corroborates earlier reports that DMBT1 protein loss in tissues is associated with poor prognosis in both colorectal^80^ and prostate^81^ cancers. A recent study also showed DMBT1 loss in a mouse model of colorectal cancer^82^, consistent with our results. However, *Dmbt1* is not a typical tumour suppressor, as results from our mouse and a previous *Dmbt1* mutant mouse that targeted exon 1^83^ showed no enhanced susceptibility to cancer. Given its physiological mitogenic function but loss during the transition to adenoma, *Dmbt1* appears to have both proto-oncogenic and tumour suppressor functions, fitting in a newly recognized class of cancer factors that tend to be lost at the protein (not genetic) level in human colorectal cancers^84^.

The main function of the IgA-DMBT1 interaction in the regulation of cell cycling appears to be homeostatic. DMBT1 is a highly glycosylated scavenger receptor with many potential ligands identified through study of its secreted forms^85^. Although it binds SIgA, it also binds other antibody structures, and our findings of the particular relevance of SIgA *in vivo* may be partially explained by the high concentrations at mucosal surfaces that no other isotype reaches, even in the context of IgA deficiency and oral treatments. Just as IgA is produced in response to microbial colonization, DMBT1 is known to be upregulated by general bacterial signals transduced via NOD2 and Toll-like Receptors^86^. DMBT1 induction by bacteria has been demonstrated in mammals^87^ and corals^88^, suggesting an evolutionarily ancient relationship, predating immunoglobulin, between this receptor and epithelial responses to microbiota. DMBT1 has previously been shown to promote proliferation of endothelial cells^89^ and was correlated with epithelial proliferation in the context of *Citrobacter rodentium* pathogenesis^90^. As part of an inflammatory response, DMBT1 is upregulated in both Crohn’s Disease and Ulcerative Colitis, and total DMBT1 knockout mice were found to be more susceptible to mouse models of colitis^91^. DMBT1 appears to be part of a stereotypical epithelial defense response to bacterial stimulation, which must be regulated in a steady state to avoid overreaction to the commensal microbiota. The present study establishes a homeostatic feedback mechanism whereby IgA inhibits the ability of DMBT1 to activate epithelial proliferation. In the absence of IgA, DMBT1 induces the proliferation of absorptive progenitors with antimicrobial properties. This mechanism may explain early microarray experiments of small intestinal samples from B cell-deficient, IgA^−/−^, and *Pigr*^−/−^ mice, which detected increased expression of genes encoding antimicrobial peptides^92,93^. While the role of IgA in shaping microbiota form and function is well established, this direct interaction with epithelial cells via DMBT1 provides a parallel function.

